# The Atlantic Forest of South America: spatiotemporal dynamics of remaining vegetation and implications for conservation

**DOI:** 10.1101/2023.09.16.558076

**Authors:** Maurício Humberto Vancine, Renata L. Muylaert, Bernardo Brandão Niebuhr, Júlia Emi de Faria Oshima, Vinicius Tonetti, Rodrigo Bernardo, Carlos De Angelo, Marcos Reis Rosa, Carlos Henrique Grohmann, Milton Cezar Ribeiro

## Abstract

The Atlantic Forest in South America (AF) is one of the world’s most diverse and threatened biodiversity hotspots. We present a comprehensive spatiotemporal analysis of 34 years of AF landscape change between 1986-2020. We analyzed landscape metrics of forest vegetation only (FV), forest plus other natural vegetation (NV), and investigated the sensitivity of metrics to linear infrastructure. Currently, remnants comprise about 23% of FV and 36% of NV, and have decreased by 2.4% and 3.6% since 1986, respectively. Linear infrastructure negatively affected large fragments (>500,000 ha) by breaking them apart. Our findings suggest that AF protection legislation adopted in mid-2005 has taken effect: between 1986-2005, there was a loss of FV and NV (3% and 3.45%) and a decrease in the number of FV and NV fragments (8.6% and 8.3%). Between 2005-2020, there was a relative recovery of FV (1 Mha; 0.6%), slight loss of NV (0.25 Mha; 0.15%) and increase in the number of FV and NV fragments (12% and 9%). Still, 97% of the vegetation fragments are small (<50 ha), with an average fragment size between 16 and 26 ha. Furthermore, 50-60% of the vegetation is <90 m from its edges, and the isolation between fragments is high (250-830 m). Alarmingly, protected areas and indigenous territories cover only 10% of the AF and are very far from any fragments (>10 km). Our work highlights the importance of legislation and landscape dynamics analysis to help monitor and keep track of AF biodiversity conservation and restoration programs in the future.

**Highlights:**

- There is 23% forest and 36% natural vegetation cover remaining in the Atlantic Forest.
- Between 1986-2020, native forest cover decreased by 2.4% and natural vegetation by 3.6%.
- Since 2005, there has been a 1 Mha increase in forest area by small fragments (1 ha).
- Roads and railways reduced by 56%-89% fragment size, especially on large fragments.
- Alarmingly, 97% of fragments are small (<50 ha) and 60% are under edge effect (<90 m).

## 1. Introduction

Habitat loss, fragmentation, and degradation caused by human-induced changes are identified as the main drivers of biodiversity loss worldwide (Chase et al., 2020). The accelerated land use conversion resulting from these changes has affected especially forest ecosystems, causing a decrease in fragment size and an increase in edge effects (Fischer et al., 2021; Hansen et al., 2020). In recent decades, tropical and subtropical regions have lost >100 million hectares (Mha) of natural forests due to anthropogenic activities (Zalles et al., 2021). Despite the large impacts, few studies presented a spatiotemporal panorama temporal long enough to describe and analyze the landscape structure dynamics, especially in the Americas, where the most diverse and threatened biodiversity hotspot in the world remains: the Atlantic Forest in South America (AF) (Sloan et al., 2014).

The AF covers almost all the coast of Brazil and portions of Paraguay and Argentina. Before European colonization, its vegetation covered over 1.6 million km^2^ (Marques et al., 2021). Due to its high environmental heterogeneity, topographic variability, and pre-historic process of formation, the AF has a high species diversity and endemism (Peres et al., 2020): it hosts more than 18,000 species of plants (Flora e Funga do Brasil, 2023) and 3,500 species of vertebrates (Figueiredo et al., 2021; Reis et al., 2016). In addition, the AF provides ecosystem services for >150 million people, such as water provisioning, hydroelectric energy generation, food production, pollination, soil protection, climate regulation, carbon storage, air quality, and cultural services (Joly et al., 2014).

The intensification of degradation arises with the Portuguese colonization and degradation of agricultural processes such as large plantation systems (sugarcane and coffee), extensive cattle production, energy demand (charcoal), fires, and urban and industrial growth (Solórzano et al., 2021). These habitat transformations have affected the biodiversity in the AF for different taxonomic groups (Püttker et al., 2020) and ecological processes, such as seed dispersal (Marjakangas et al., 2020), carbon storage (de Lima et al., 2020), pollination (Varassin et al., 2021), and top-down regulation through top predators (Paviolo et al. 2016). In addition, other processes pose risks to the remaining landscapes within the AF, such as defaunation (Galetti et al., 2021) and climate change (Vale et al., 2021).

Despite the recent changes on the AF, few studies have analyzed the landscape structure in a space-time context on large time scales. In the most comprehensive study to our knowledge, Ribeiro et al. (2009) showed that only 11-16% of the forest cover remained in 2005, 83% of which was concentrated on isolated fragments smaller than 50 ha, and half of all forests were <100 m from their edges. After that, Tabarelli et al. (2010) and Ribeiro et al. (2011) showed a large proportion of forests remained in high elevations (>1600 m). Based on finer scale satellite data (5 m-spatial resolution), Rezende et al. (2018) estimated 28% of remaining AF vegetation. In more recent studies, using data from MapBiomas (Souza et al., 2020), Bicudo da Silva et al. (2020) showed that landscape composition did not change between 1985-2018, and that the loss in areas of montane vegetation was smaller than at lower elevations. Rosa et al. (2021) showed that the relative temporal stability of AF native forest cover (28 Mha) in recent years, was in fact due to the loss of old-growth native forests in flatter terrains, and the growth of young forests in marginal agricultural areas, resulting in increased isolation.

Despite these studies, there is a demand for refined data to understand how landscape structure varied over time in AF. Currently, Brazilian initiatives such as MapBiomas have been mapping land use and land cover (LULC) change with wide thematic coverage, high spatiotemporal resolution, and standardized classification (Souza et al., 2020). This allows for the calculation and comparison of landscape metrics for large territorial extensions and time periods to understand the landscape dynamics of entire domains (Bicudo da Silva et al. 2020; Rosa et al. 2021). In addition, the AF has a high density of linear infrastructure since it hosts a high (and increasing) human population. This severely impacts natural vegetation and biodiversity and must be considered in landscape structure analyses (Cassimiro et al., 2023).

Here, we analyzed the spatiotemporal dynamics of the landscape structure of vegetation in the AF every five years from 1986-2020. To accomplish this large-scale evaluation, we used a wide delimitation of Atlantic Forest, including Brazil, Argentina, and Paraguay. We accounted for forest vegetation types only (FV) and forest plus other natural vegetation types (NV) and quantified the effect of linear infrastructure on the AF landscape metrics. To understand the spatiotemporal vegetation dynamics, we calculated the following landscape metrics for all FV and NV fragments in the AF domain: fragment size, number of fragments, fragment temporal dynamic, habitat amount, edge area, isolation, functional connectivity, and distance from protected areas (PA) and indigenous territories (IT). These metrics were generated through an approach that allows an ecological interpretation of the influence of the landscape structure on organisms, by accounting for species mobility, gap-crossing abilities, and sensitivity to edge effect (Riva and Nielsen, 2020).

## 2. Methods

### 2.1 Study region

AF extends from 3°S to 33°S, and from 35°W to 58°W with about 163 Mha, covering large coastal and inland portions of Brazil, Argentina, and Paraguay (Marques et al., 2021) (Fig. S1a). Due to this extension, the AF boundaries create important ecotones with other vegetation domains such as Cerrado, Caatinga, Chaco and Pampa (Marques et al., 2021). The vegetation from AF is a complex mosaic mainly composed of five vegetation types— Dense Ombrophilous, Open Ombrophilous, Mixed Ombrophilous, Semideciduous Seasonal, and Deciduous Seasonal (Joly et al., 2014). Additionally, the AF also includes mangroves and coastal scrub vegetation (Marques et al., 2021). Besides, there are many marginal habitats such as altitude grasslands (*campos rupestres* and *campos de altitude*), oceanic islands, beaches, rocky shores, dunes, marshes, inland swamps, and mountain forest (*brejos de altitude*) in the Northeast region (Scarano, 2002). Therefore, we used an integrative delimitation adapted from Muylaert et al. (2018), which encompasses the main proposed delimitations across several associated ecosystems. This delimitation was produced by overlapping available AF delimitations (Table S1 and Fig. S1[b-e]) and adjusting the delimitation in the Eastern coastal areas using the Brazilian territorial delimitation from IBGE (https://www.ibge.gov.br) for 2021. This step ensures that areas of coastal vegetation such as mangroves, dunes, and wooded sandbank/sandy coastal plain vegetation (hereafter *restinga*) (Scarano, 2002) are better represented. The final delimitation has a total area of 162,742,129 ha, distributed within 3653 municipalities from 18 Brazilian states (93.1%), 70 municipalities of one province in Argentina (1.6%), and 127 municipalities from 11 departments in Paraguay (5.3%) (Fig. S1a).

### 2.2 Mapping

We compiled LULC maps for Brazil, Argentina, and Paraguay from MapBiomas Brazil collection 7 (https://mapbiomas.org/) and MapBiomas Bosque Atlántico collection 2 (https://bosqueatlantico.mapbiomas.org/) (Souza et al., 2020). These datasets reconstruct annual LULC information at 30-m spatial resolution from 1985 to 2021, based on a pixel-based random forest classifier of Landsat satellite images using Google Earth Engine, with AF general accuracy of 89.8% (Souza et al., 2020). We used the interval beginning in 1986 and ending in 2020. We excluded the years 1985 and 2021, as there was no validation for the previous and subsequent year, respectively. Furthermore, we defined two vegetation classes for analysis: only forest vegetation types or “Forest Vegetation” (FV) and both forest and other natural vegetation types or “Natural Vegetation” (NV) (Table S2), for every fifth year between 1986 and 2020 (Fig. S2a-h).

We used roads and railways to trim their overlapping FV and NV (henceforth called “trimmed” and “not trimmed” scenarios). This procedure enabled us to avoid overestimating large fragments of vegetation and check the metrics’ sensitivity to linear infrastructure, since these structures decrease landscape connectivity and threaten multiple taxonomic groups (Cassimiro et al., 2023). Thus, we analyzed four vegetation maps: “FV not trimmed”, “FV trimmed”, “NV not trimmed”, and “NV trimmed”. Further, we analyzed the overlap between FV and NV fragments with Protected Areas (PA) and Indigenous Territories (IT). Details of road, railway, PA, and IT maps are presented in the Data section in the Supplementary Material. All geospatial datasets were rasterized and warped to 30 m-spatial resolution (112663 × 83307 ≈ 9.4 billion cells) using the Albers Conical Equal Area Brazil (SIRGAS 2000) projection (https://spatialreference.org/ref/sr-org/albers-conical-equal-area-brazil-sirgas-2000/). International map displays were generated using Natural Earth (1:10,000,000) data and QGIS 3.22 LTR (QGIS Development Team, 2023).

### 2.3 Landscape metrics

All landscape metrics were processed in GRASS GIS 8.2.1 (Neteler et al., 2012) through the R 4.3.0 (R Core Team, 2023), using the *rgrass* package (Bivand, 2022). We calculated six landscape metrics: number of fragments, fragment size, edge area, isolation, functional connectivity, and distance from PA and IT (Table S3 and Figure S13). The number of fragments and fragment size allowed us to account for the number and area of remaining vegetation fragments for different size classes (Table S3). Fragments were defined using the eight-neighbor rule (Queen’s case), which defines areas connected to pixels in eight directions (Turner and Gardner, 2015). We also examined the area and number of fragments that appeared and disappeared throughout time, and the areas of increase, reduction, and stability of fragments that remained in the landscape (Table S3) (Rosa et al., 2021). Edge area was calculated for different edge depths (distance from the edge of the fragment) (Table S3), allowing us to assess the amount and percentage of forest area subjected to edge effects (Harper and Macdonald, 2011).

Two metrics of functional connectivity were computed for different gap-crossing distances (species’ capacities to cross the non-habitat) (Table S3). First, we calculated the sum of the areas of all fragments closer than the gap-crossing distance, which can be interpreted as the functional available area of each clump of fragments (Awade and Metzger, 2008). Second, we computed the expected cluster size as the mean fragment clump size, and then compared it with the highest cluster size in the entire study region. Isolation was calculated using an index adapted from the ‘‘Empty Space Function” (Dale and Fortin, 2014), similar to Ribeiro et al. (2009): we computed a Euclidean distance map from all the fragments, extracted its values and calculated the mean. We repeated this process by removing different-sized fragments in several steps (see Table S3 for classes of distances), and then created new Euclidean distance maps to recompute the mean distance values.

These values represented the isolation of fragments while also providing insights about the importance of the smaller fragments (*stepping stones*) (Diniz et al., 2021). We calculated the amount of FV, NV, and vegetation classes (see Table S2) covered by PA and IT, and the shortest Euclidean distance from each FV and NV pixel to these areas (see Table S3 for classes of distances).

## 3. Results

### 3.1 Number of fragments and fragment size distribution

Roads and railways greatly impacted the large-sized fragments, depending on the year and the scenario considered. These effects were mainly reflected in vegetation fragments larger than 500,000 hectares, for which the maximum fragment size decreased by 56%-89% (Fig. 1, Fig. S3, and Table S4). By accounting for linear infrastructure, the >1 Mha fragment size class ceased to exist for FV for all years and was heavily reduced for NV, and the total area and number of fragments increased for fragments of all size classes <500,000 ha for FV and NV (Fig. 1, Fig. 2, and Fig. S4). Despite this effect for large fragments, our results showed no difference between the scenarios “trimmed” and “not trimmed” for other landscape metrics. Therefore, we chose to demonstrate the results with the linear infrastructure effect (trimmed scenario) in the main text and present the additional results in the Supplementary Material.

**Figure 1.**
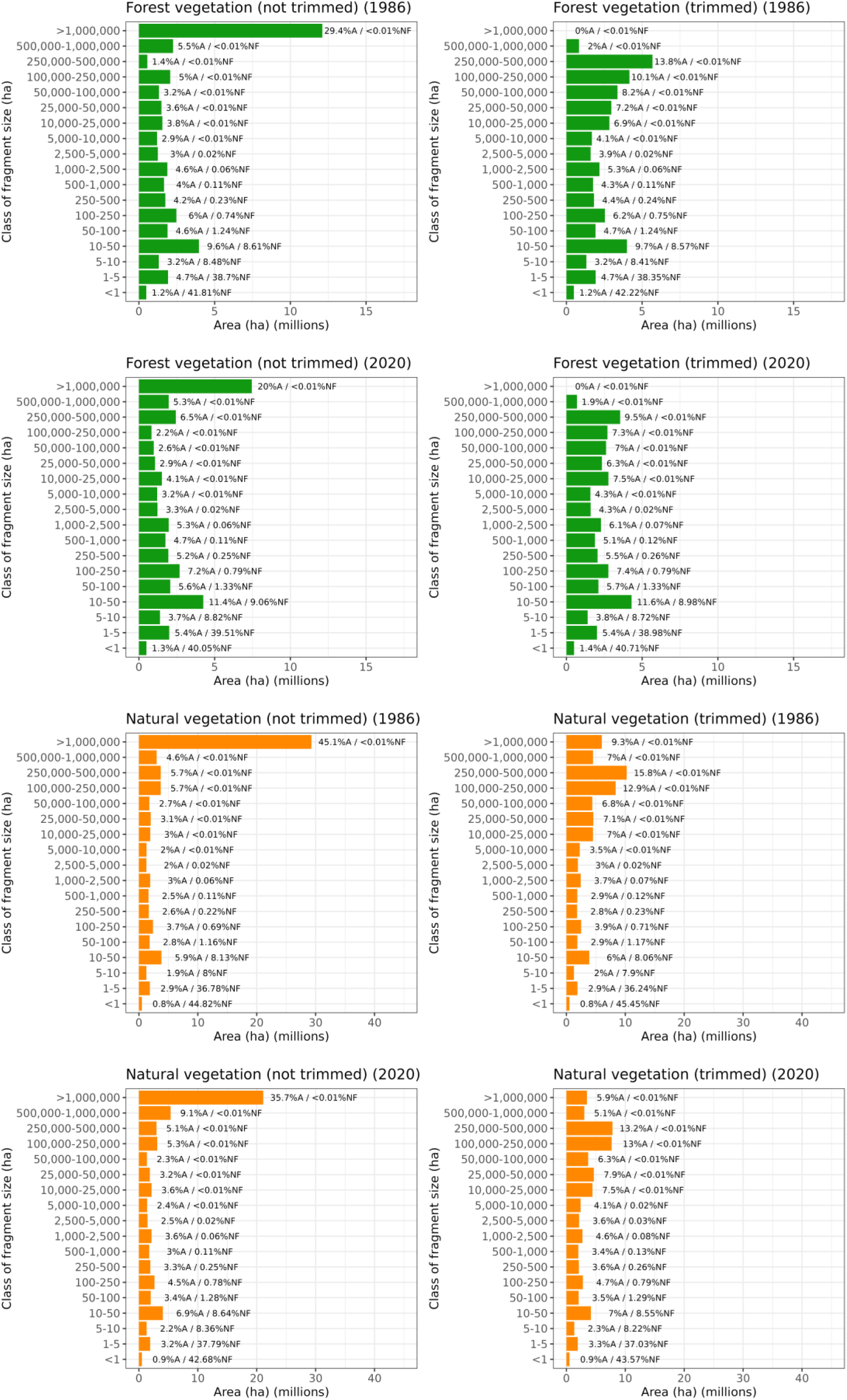
Distribution of FV and NV fragment sizes across the AF (1986 and 2020), trimmed and not trimmed by linear infrastructure. %A: percentage of the total area; %NF: percentage of the number of fragments. See Fig. 3S for other years (1990-2015). Please note the difference scales in the x-axis between the FV and NV plots.

**Figure 2.**
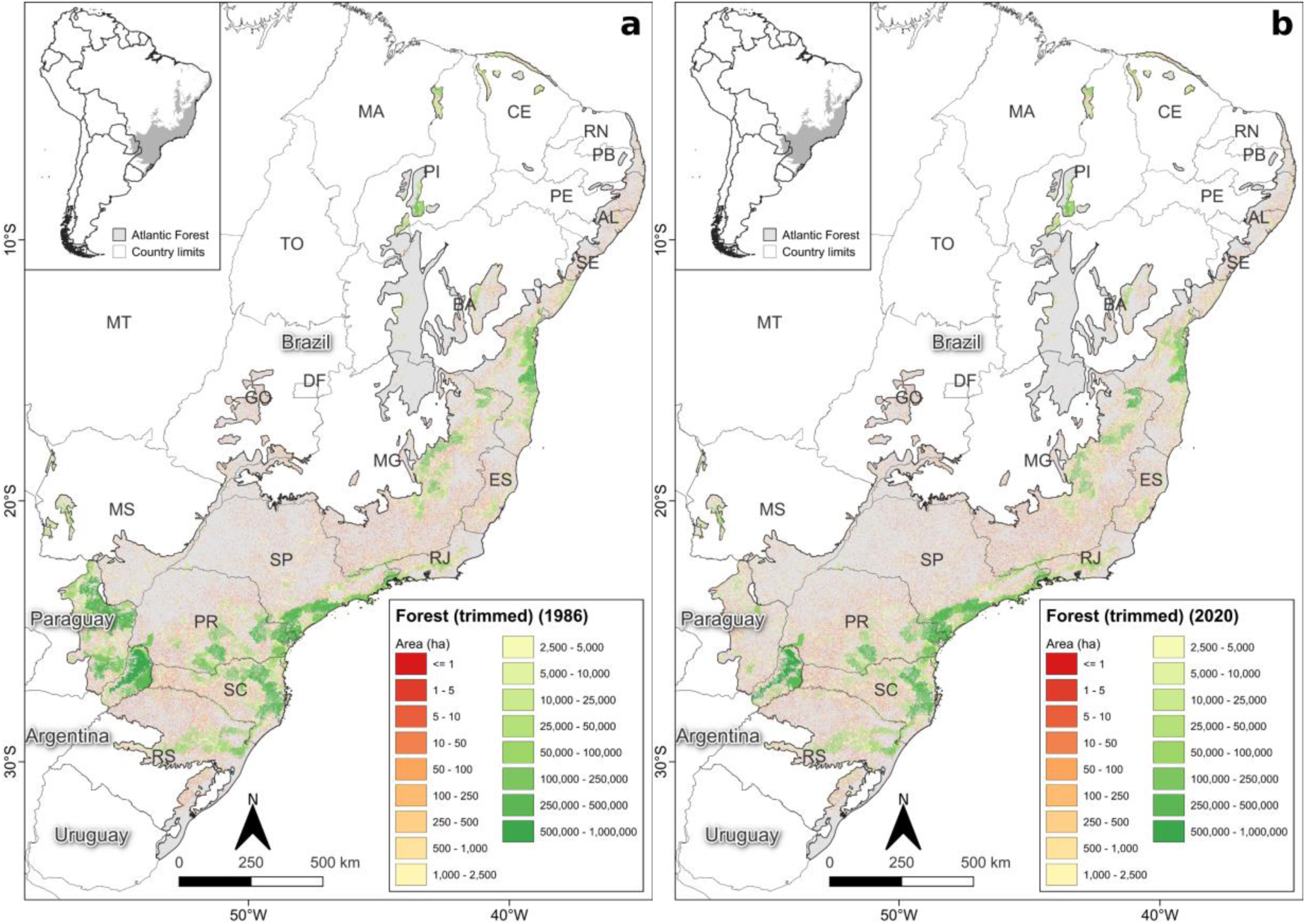
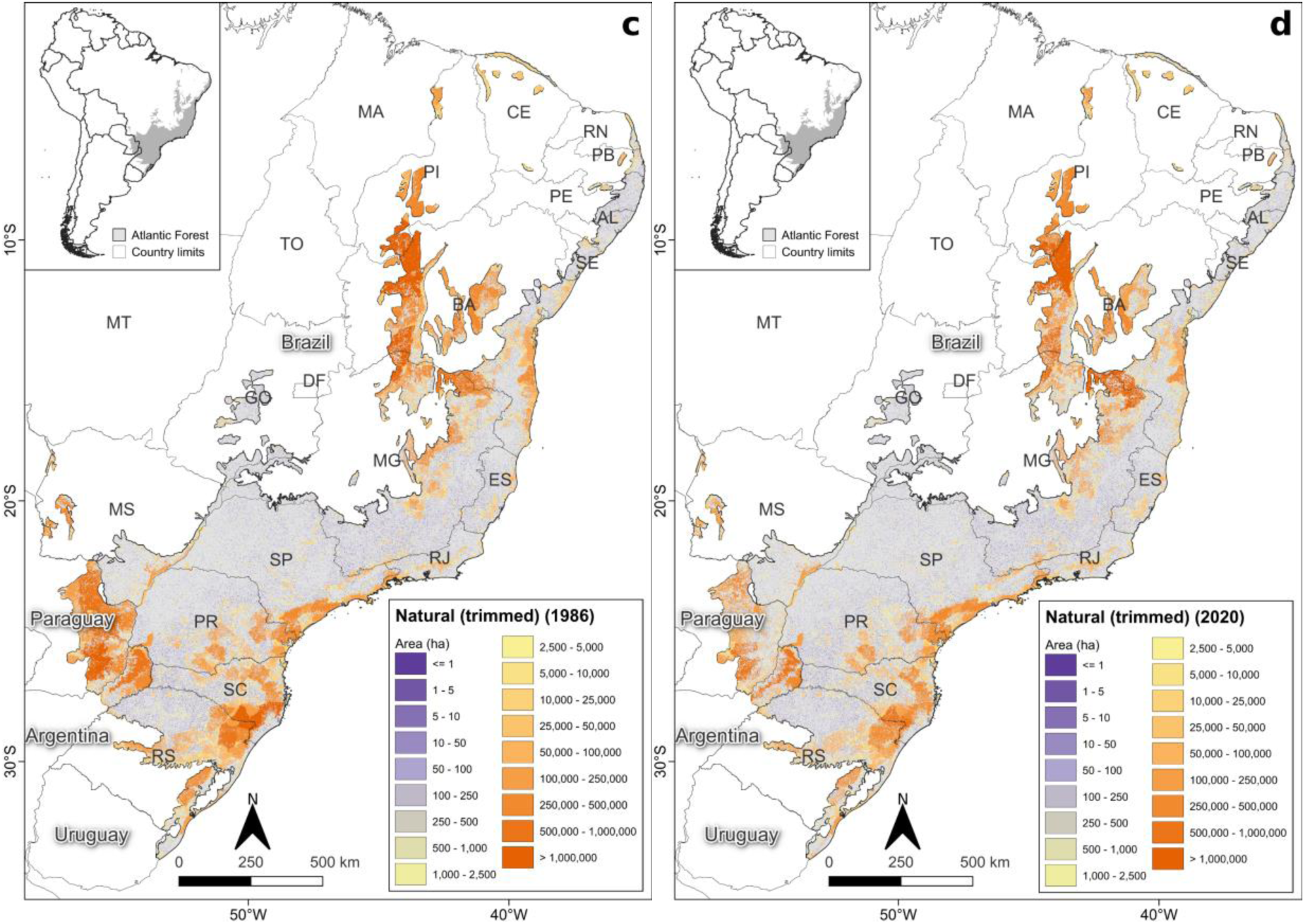
Fragment area for FV in 1986 (a) and 2020 (b), and for NV in 1986 (c) and 2020 (d), trimmed for the entire AF.

For the trimmed scenario, about 97% of the fragments have an area of less than 50 ha, with 0.3% of variation over the years. However, between 1986 and 2020 the total area increased from 18.8% to 22.1% for FV and from 11.6% to 13.4% for NV (Fig. 1 and Fig. S3). For fragments between 50 ha and 25,000 ha, the proportion of the total number of fragments is low (2.5%), varying for FV from 2.44% in 1986 to 2.66% in 2020, with a maximum value of 2.76 % in 2005; and for NV with 2.34% in 1986 to 2.61% in 2020, and a maximum of 2.66% in 2005. However, total area increased from 1986 to 2020, going from 39.8% to 45.9% for FV, but very similar since 2005 (45.1%); and for NV, from 29.6% to 35.1% (Fig. 1 and Fig. S3). For the last category of fragment area, above 25,000 ha, we found a very small proportion of number of fragments (0.001%), with values falling from 0.0081% to 0.0058% for FV, and from 0.0125% to 0.0116% for NV, between 1986 and 2020. Total area values for FV fragments in these categories fell from 41.4% to 32% and for NV from 58.7% to 51.5%, between 1986 and 2020 (Fig. 1 and Fig. S3).

In 1986, the largest FV fragments were localized in the coast of Bahia (*South of Bahia*—*cabruca region*), São Paulo, Paraná, and Santa Catarina states (*Serra do Mar region*), and inland areas of Paraná, Santa Catarina and Rio Grande do Sul states in Brazil. For the same period, there were large FV fragments in the Misiones region in Argentina and the east portion of Paraguay (Fig. 2a). We observed the same for NV, with additions of huge fragments in portions of Bahia, Minas Gerais, and Piauí states, mainly in the regions named *São Francisco* and *Brejos Nordestinos* (see these region concepts in Ribeiro et al., 2009) (Fig. 2c). In 2020, these same regions concentrated the largest fragments of FV and NV, but with a decrease in the area of these fragments (Fig. 2[b-d]). The exception was Paraguay, where there was a vast deforestation process, mainly for FV (Fig. 2[c-d]). Our results also show a large effect of roads and railways on maximum fragment size, for the same regions (compare Fig. 2[a-d] with Fig. S4[a-d]).

The spatial-temporal analysis revealed a turning point for the AF landscape structure in 2005. For the first period (1986-2005), the number of fragments decreased by 8.6% for FV and 8.3% for NV; for the second one (2005-2020), it increased to 11.9% for FV and 9% for NV (Fig. 3a). From 2010 onwards, the number of FV and NV fragments tended to become more like each other (Fig. 3a). The average fragment size in the first period for FV dropped by 3.5% (18.5 to 17.9 ha) and remained stable for NV, dropping 0.3% (28.5 to 28.4%). In the second period, the FV had a great drop of 8.2% for FV (17.9 to 16.4 ha) and 8.6% for NV (28.4 to 25.9 ha) (Fig. 3b).

**Figure 3.**
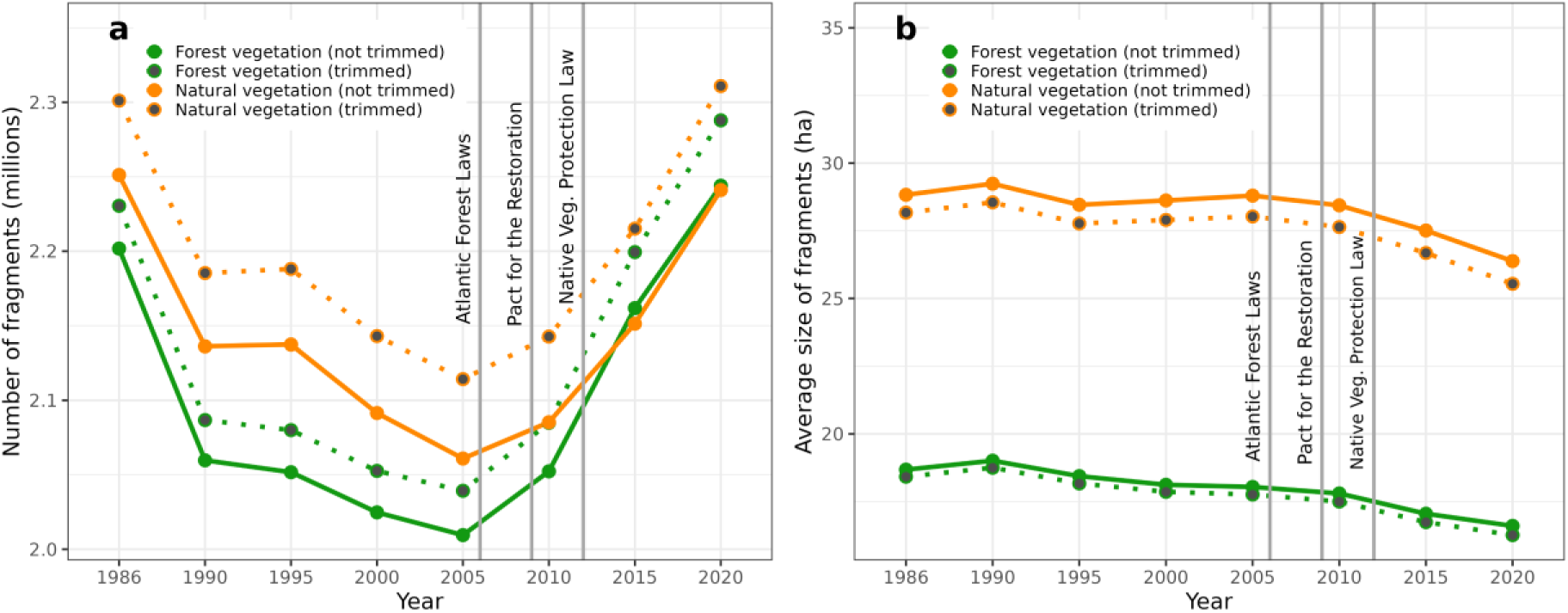
Distribution of number of fragments (a) and average fragment sizes (b) of FV and NV across the AF from 1986 to 2020, trimmed and not trimmed by roads and railways. The gray lines represent when the legislation and restoration programs were created (See Discussion for details).

The temporal dynamics of the landscape (Table S3) from 1986 to 2005 revealed a reduction in the total area of 4.78 Mha of FV (3%) and 5.56 Mha of NV (3.4%) (Table S5). However, between 2005 and 2020, there was an increase of 985,000 ha of FV (0.6%) and a small decrease of 240,000 ha of NV (0.15%). Considering the balance of fragments gained and lost, in the first period there was a sharp drop in the number of fragments for FV (242,000) and NV (227,000), but in the second period there was an increase for FV (380,000) and NV (310,000). Between 1986-2005, the average size of lost FV and NV fragments (1.2 to 1.35 ha) was greater than the size of restored fragments (1.08 to 1.14 ha); between 2005-2020, this pattern reversed, with the average size of fragments lost was smaller (0.94 to 0.97 ha) than that of fragments gained (1.03 to 1.08 ha).

### 3.2 Forest and natural vegetation cover

The proportion of the Atlantic Forest domain covered by forests and natural vegetation decreased in the past 34 years, from 25.26% (41.1 Mha) to 22.86% (37.2 Mha) for FV and from 39.86% (64.8 Mha) to 36.27% (59 Mha) for NV (Fig. 4 and Table S4). For the entire period, in Brazil the percentages decreased from 22.85% (34.6 Mha) to 22.27% (33.7 Mha) for FV and from 37.34% (56.5 Mha) to 35.25% (53.4 Mha) for NV, with a stable proportion since 2005 (Fig. S5). NV was mainly composed of savannas, grasslands, and wetlands, besides the forest formations. In Argentina, the loss of vegetation cover was proportionately larger, from 67.38% (1.8 Mha) to 56.9% (1.52 Mha) for FV, and 67.99% (1.82 Mha) to 57.34% (1.53 Mha) for NV, showing an increase in the rate of deforestation in the last five years (Fig. S5). In Paraguay, the loss of vegetation cover was higher than in the other countries, dropping from 54.57% (4.7 Mha) to 22.85% (2 Mha) for FV, and 75.26% (6.5 Mha) to 47.74% (4.1 Mha) for NV (Fig. S5), but it has maintained its remnants since 2005.

**Figure 4.**
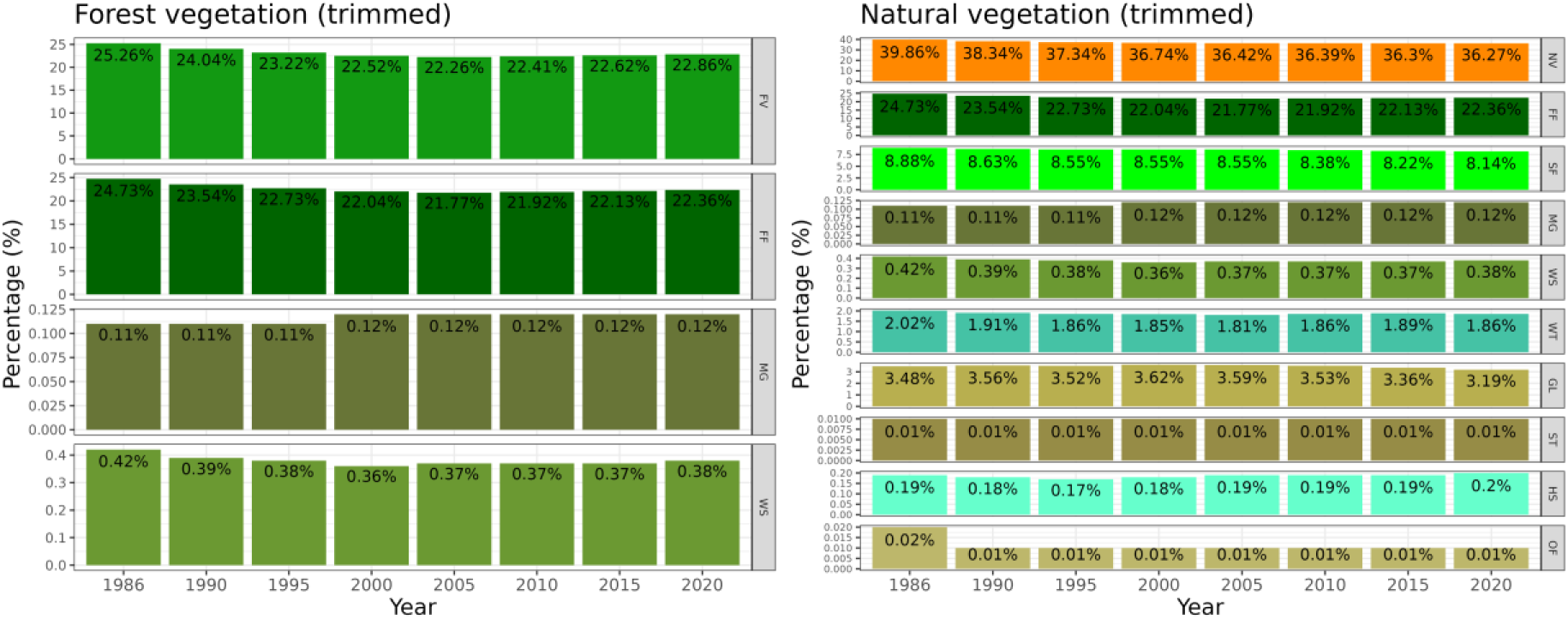
Vegetation cover for FV and NV through the years, trimmed. Abbreviations in Table S2.

Beyond presenting the percentages for the integrative delimitation (Fig. S1a), we also present results for five other delimitations (Table S6). The results for 2020 (Fig. S1[b-f]) varied for FV from 23.15% (31.6 Mha) for the delimitation of Da Silva and Casteleti (2003) trimmed up to 26.70% (32.2 Mha) for the delimitation of Dinerstein et al. (2017) disregarding the effect of roads and railways. The same occurred for NV, ranging from 31.45% (34.8 Mha) for the IBGE (2019) trimmed to 35.98% (46.3 Mha) for the delimitation of the Atlantic Forest Law (2006) not trimmed (Table S6).

### 3.3 Core and edge area

The percentage of FV and NV remaining less than 90 m from the edge increased over time, going from 52% to 59% for FV and 42% to 48% for NV, as well as the percentage less than 240 m, from 76% to 82% for FV and 66% to 72% for NV (Fig. 5[a-b]). Conversely, the amount of FV and NV more than 500 m from any edge decreases, from 12% to 9% and from 20% to 15%, respectively. The maximum edge distances for FV and NV were quite different, being around 11 km for the FV, and 32 km for the latter, showing that NV creates large core areas (Fig. 5[a-b]). From 90 m onwards, there is an inversion in the edge percentage over time: <90 m, there is a gradual increase in the percentage between 1986 and 2020; >90 m, the percentage of vegetation starts to decrease, showing the conversion of fragment core areas to edge areas (Fig. 5[c-d]).

**Figure 5.**
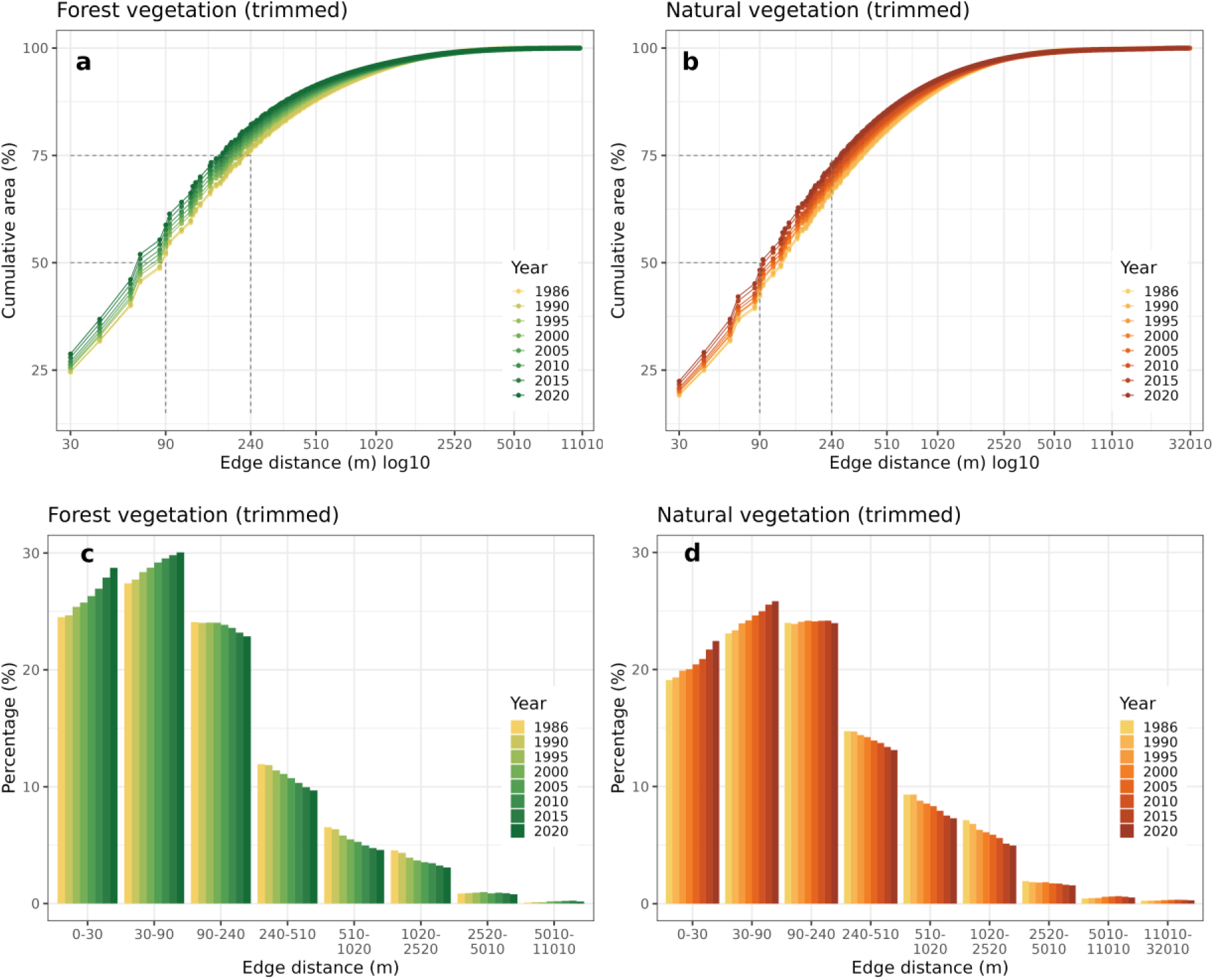
Cumulative (a and b) and per class (c and d) area under edge effect at different depths for the FV and NV remaining in AF trimmed.

### 3.4 Functional connectivity

Considering the functional connectivity for species that cannot cross areas of non-habitat (i.e., gap-crossing equals 0 m), the average functionally connected area for FV decreased 11.7% (18.42 to 16.26 ha) (Fig. 6a), and 9.3% for NV (28.2 to 25.5 ha) between 1986 and 2020 (Fig. 6b). The same pattern occurs for 60 m of gap-crossing for both types of vegetation. However, for gap-crossing values between 120 and 180 m, functional connectivity decreased until 2005 and then increased. For values above 240 m, functional connectivity also decreased until 2005, then increased, with its value in 2020 greater than in 1986. The functional connectivity of the NV was always higher in numerical terms for the same years, but they followed the same patterns of annual trends and gap-crossing of the FV.

**Figure 6.**
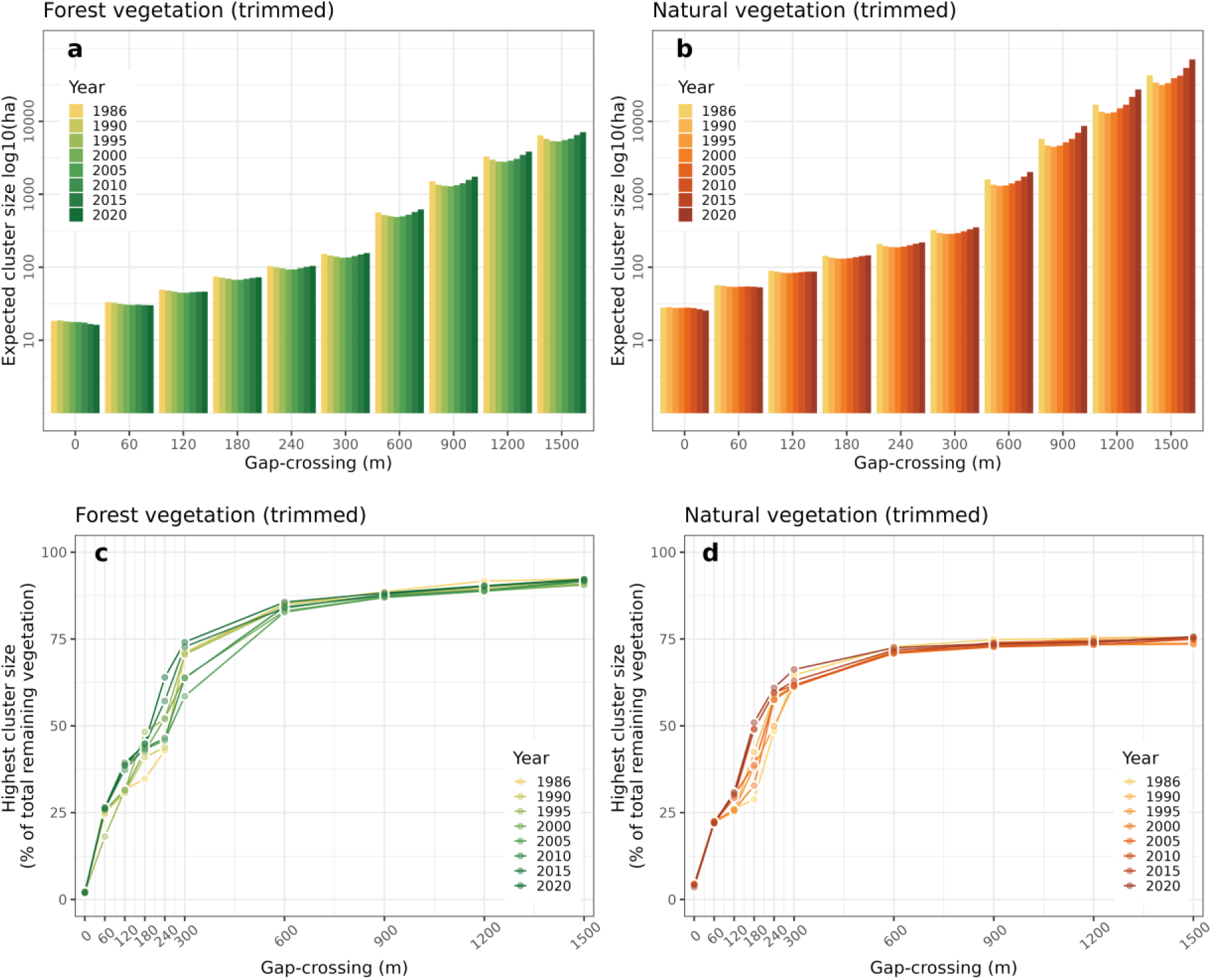
Expected cluster size (a-b) (average functional size; ha) of functionally connected fragments of FV and NV for different functional distance values (meters) for the AF trimmed by roads and railways. Highest functionally connected vegetation cluster (c-d) (% of total remaining of FV and NV) estimated across varying functional distances (meters) for the AF.

When we analyzed the highest functionally connected vegetation cluster, we noticed similarities in the pattern of the curves generated for FV and NV: both curves started with a low relation between the highest functionally connected FV and NV cluster (about 2% for FV and 4% for NV for all years), indicating low connectivity when we do not consider a value of gap-crossing (Fig. 6[c-d]). Despite that, around 600 m of gap-crossing, both reached their respective horizontal asymptotes, which were about 90% for FV and 75% for NV for every year, with a significant divergence between years between 60 and 600 m cross-level difference values, ranging from 18% to 75%. For the not trimmed scenario, there was a difference in the highest functionally connected initial percentages due to the different values of the largest fragments over the years (Fig. S7[c-d]).

### 3.5 Mean isolation

Small fragments were key to reducing isolation in all analyzed scenarios. For example, when we disregard fragments <50 ha, the isolation increases to 79-83% for FV and 78-85% NV (Fig. 7[a-b]). Furthermore, isolation was highly reduced in 60-85% when considering NV also for all temporal scenarios and road and rail effects analyzed (Fig. 7[a-b], Fig. S8[a-b]). In 1986, the mean isolation for the entire AF region was 773 m for FV and 273 m for NV. The isolation reached its maximum values in 1995, with values of 949 m and 291 m for FV and NV, respectively. After that, the isolation had a slow decrease until 2015, going to 902 m for FV and 266 m for FV; and more recently, it fell to 832 m for FV and 253 m for NV in 2020 (Fig. 7[a-b]). When we disregarded fragments smaller than the size classes defined in Table S3, there was a significant increase in isolation, a gradual increase for each size that we disregarded, varying to 4-22 km for FV and 1-12 km for NV. In general, isolation peaks for each class fragment size disregarded occurred between 1990 and 2000 for FV and NV, which after these dates began to decrease, reaching the lowest values of the historical series in 2020, both for FV and for NV.

**Figure 7.**
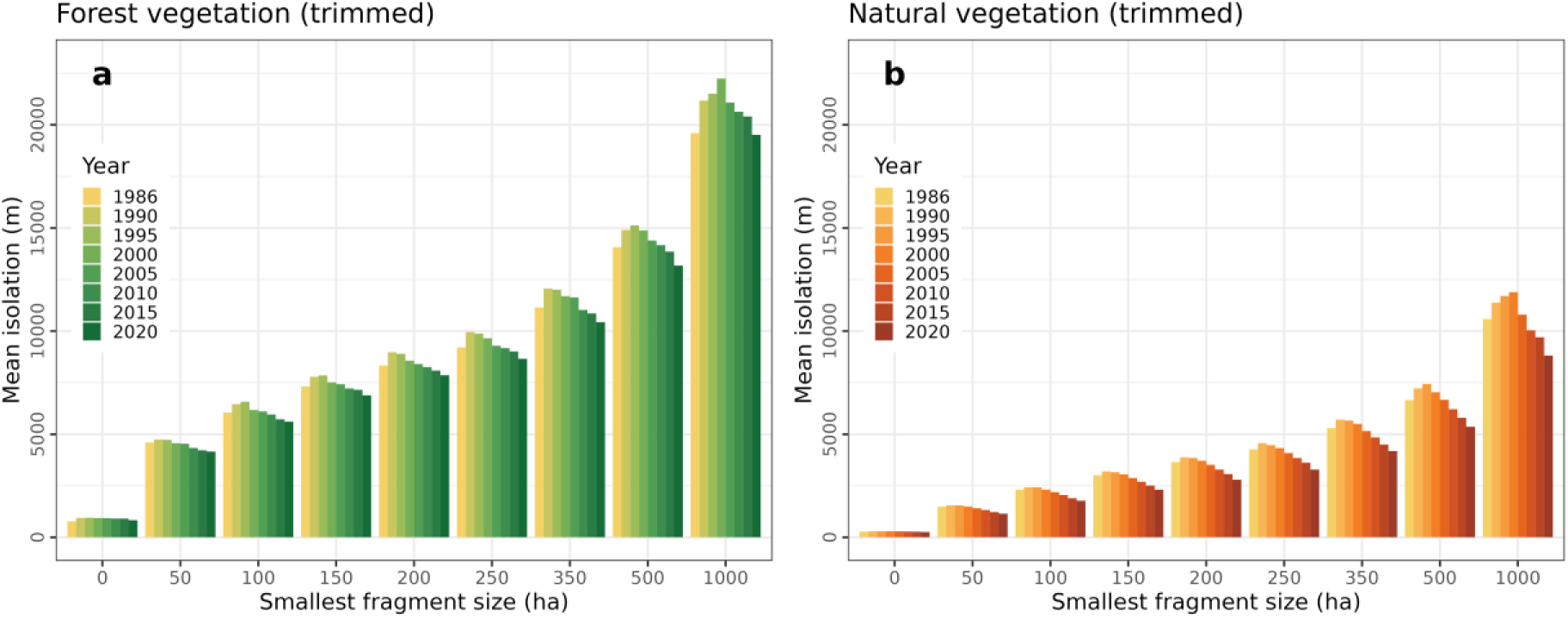
Influence of the smallest fragment size (ha) on the isolation (m) between fragments of FV and NV trimmed for the AF. Smallest fragments size: 0 ha (all fragments), 50 ha, 100 ha, 150 ha, 200 ha, 250 ha, 350 ha, 500 ha, and 1000 ha.

### 3.6 Protected areas and indigenous territories distance

Protected areas (PA) covered 4.6 Mha (2.84%) and indigenous territories (IT) covered 1.3 Mha (0.81%) of the AF limit. These values represent 12.4% and 7.8% of the total FV and NV area for PA, and 3.6% and 2.2% of the total FV and NV area for IT in 2020. However, only 3.1 Mha (8.4%) of FV and 4.1 Mha (7%) of NV remaining overlaps with PA (Fig. 8[a-b]), and only 0.56 Mha (1.5%) of FV and 0.76 Mha (1.3%) of NV overlaps with IT (Fig. 8[c-d]), since other types of land cover occur within PA and IT. Only 2.7% of FV and 2.2% of NV are within 1 km of PA and 0.8% of FV and 0.7% of NV to IT. For vegetation within 10 km, there are 23.4% of FV and 19.2% of NV of PA, and 9.5% of FV and 8.7% of NV of IT (Fig. 8). On the other hand, 68.2% of the FV and 73.9% of NV are over 10 km away from PA, and 89% of the FV and 90.2% of NV are over 10 km away from IT, demonstrating the lack of protection for these fragments of remaining vegetation (Fig. 8).

**Figure 8.**
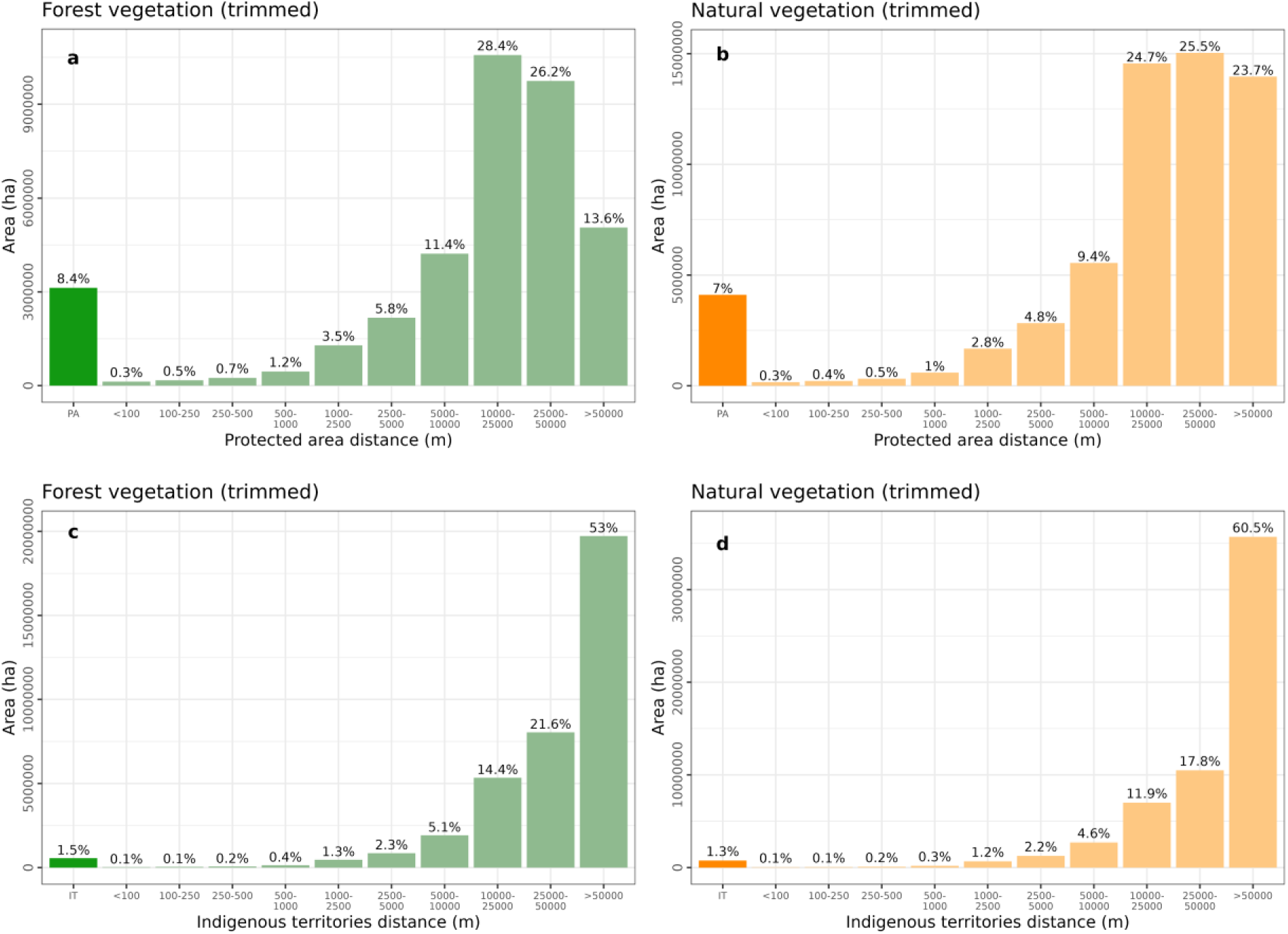
Remaining vegetation of AF remnants (area and percentage) and their distance (meters) from protected areas (PA; a – FV and b – NV) and indigenous territories (IT; c – FV and d – NV) per class, trimmed.

The vegetation class with the largest area overlap is the Forest formation, with 3 Mha (8.2%) for PA and 536,000 ha (1.5%) for IT (Fig. S10). The class with the largest cover under protection is the *restinga* with 133,000 ha (21.6%), Mangrove with 23,000 ha (11.9%), and Herbaceous sandbank vegetation with 31,000 ha (9.6%) (Fig. S10b). For IT, the class with the most cover proportion is the Other non-forest formations 1,400 ha (8.6%), followed by *restinga* with 22,000 ha (3.6%), and Mangrove with 4,600 ha (2.4%) (Fig. S10d). The Savanna formation class is the smallest cover protection with only 507,000 ha (3.8%) in PA (Fig. S10b) and 120,000 ha (0.9%) in IT (Fig. S10d), whose total area is second in terms of total area, 13 Mha (22.5%).

## 4. Discussion

### 4.1 Main results

Our results showed great changes in the spatial-temporal dynamics of the landscape structure for the entire AF, emphasizing the likely great importance of environmental legislation for its conservation. These changes varied depending on the vegetation types and roads and railways scenarios considered. For example, fragment size was highly sensitive to linear structures, especially for large fragments of FV and NV (>500,000 ha), but other metrics were not affected as percentage, connectivity, or isolation. In 34-year period, there was a high loss rate for both FV and NV, with apparent differences in trends between Brazil, Argentina, and Paraguay. Our findings revealed yet a turning point for the Atlantic Forest after 2005. In the first period (1986-2005), the number of vegetation fragments and forest cover decreased, following the strong trend of forest loss from the previous years. In the second period (2005-2020), the number of vegetation fragments increased, and the forest cover was more stable. Nevertheless, more than 97% of fragments are smaller than 50 ha, and mean fragment size decreased being currently equal to 16 ha for FV and 26 ha for NV. Besides, there was an increase in the amount of fragment closest to the <90 m from edge (50-60%), although the insulation is decreasing, reaching values equal to 830 m for FV and 250 m for NV, with the connectivity increasing.

These results bring a new panorama to the remaining AF vegetation, mainly because our analysis was more spatially and temporally comprehensive, integrated different types of vegetation, and considered for the first time a wide distribution of AF, including Argentina and Paraguay. Although legislation and restoration actions appear to be positively affecting vegetation restoration processes, there is still a scenario of intense habitat fragmentation. Allied to this, NV played a fundamental role in increasing the connectivity of the vegetation remnants, despite having been little favored by protection areas. Our results also showed that although there is an overlap of about 10% of the vegetation with PA and IT, a large part of the vegetation is located far from these areas (>10 km) and given the great anthropic concentration in the AF and recent urban expansion, a portion significant amount of vegetation could be impacted in the recent future.

### 4.2 Number and fragment size distribution

Our analyses of temporal dynamics demonstrated that between 2005-2020 there was an increase in habitat amount, number of fragments, and higher mean fragment gain for FV. This corroborates that there was a regeneration process since new fragments appeared, which confirms the results found by Rosa et al. (2021) and Dias et al. (2023), who highlighted the replacement of older vegetation by younger vegetation in the AF. However, this replacement can lead to the loss of quality of habitat fragments, altering landscape features, and affecting vital ecological processes and ecosystem functioning, such as carbon cycling (Piffer et al., 2022) and vegetation structure (Faria et al., 2023). The effect of roads and railways was more pronounced in FV than in NV, due to their greater density in large forest fragments located in Serra do Mar, southern Bahia in Brazil, and in the region of Misiones in Argentina. Roads and railways have a huge impact on biodiversity, modifying the movement pattern, reducing connectivity, and causing roadkill, which leads to population declines and local extinction (Cassimiro et al., 2023; Martinez Pardo et al., 2023).

When analyzing the distribution of fragment sizes, we noticed a reduction in the number and percentage in relation to the total area of the remnants for fragments >1 Mha of FV and NV between 1986 and 2020 and a clear increase in fragments <50 ha. These results showed a worrying pattern, much worse than Ribeiro et al. (2009), and can be explained by the increase in mapping quality, with MapBiomas standardized approaches for mapping vegetation fragments (including fragments <3 ha) that are considered secondary vegetation in detail (Rosa et al., 2021). The increase in the proportion of smaller fragments has a direct impact on the maintenance of species diversity and population size of multiple taxonomic groups. Several works have estimated fragment size and habitat amount thresholds for assemblage diversity in AF, such as terrestrial mammals (Magioli et al., 2015), bats (Muylaert et al., 2016), birds (Barbosa et al., 2017), and multiple groups (Banks-Leite et al., 2014). However, since 97% of the fragments are <50 ha in AF, the general scenario is already well under the thresholds that are known to affect biodiversity composition. Then, approaches for conservation should be comprehensive and focus on single large *and* several small (SLASS) fragments (Szangolies et al., 2022). The SLASS approach can be more beneficial for conserving the AF biodiversity than choosing a unique type of conservation approach (SLOSS debate, see Fahrig et al. 2021).

### 4.3 Forest and natural vegetation cover

Determining how much of the AF vegetation cover is left has always been a complex task. While we found 22.86% for FV and 36.27% for NV using an integrative AF delimitation in 2020, for the same year, the vegetation cover values varied from 23.15% to 35.98% for different delimitations and vegetation types. Here, we highlighted the use of MapBiomas mapping version 7, with image standards and classification methods, making the comparison of annual maps possible due to the decrease in random error between them (Souza et al., 2020). Over the years, several studies have shown values ranging from 8% to 28% for different years, AF delimitation, and vegetation types (Bicudo da Silva et al., 2020; Da Silva and Casteleti, 2003; Rezende et al., 2018; Ribeiro et al., 2009). These estimates vary according to mapping resolution, size of vegetation fragments, types of vegetation (forest or non-forest), vegetation quality (primary or secondary forest), and AF delimitation (Ribeiro et al., 2009). Therefore, we emphasize that these percentage values must be used with awareness of their calculation specificities and limited comparability. Due to their simplification and high variability intrinsic to each source, their values must be presented to meet detailed and well-defined objectives, meeting specific criteria.

The vegetation cover showed a considerable decrease over time, mainly between 1986 and 2005. After 2005, the percentage of vegetation stabilized or increased, mainly in Brazil and for FV. These effects can be related to nature conservation laws, which were initiated almost in the same period in Brazil (Atlantic Forest Law in 2006, and Native Vegetation Protection Law in 2012), Argentina (Forest Law in 2007), and Paraguay (The Zero Deforestation Law in 2004) (Silva et al., 2017; Dam et al., 2019). In Brazil, specific conservation laws were established from 1998 on (Fauna Protection in 1988, and National System of Conservation Units (SNUC) in 2000), and more recently the 2012 legislation created the Rural Environmental Registry (CAR), which requires environmental information from private rural properties. CAR can be a fundamental tool to direct vegetation restoration efforts through legal reserves (LR) and permanent preservation areas (PPA) (da Silva et al., 2023). Since 2009, the Pact for the Restoration of the Atlantic Forest (https://pactomataatlantica.org.br) has been encouraging the restoration with the goal to restore 15 Mha by 2050 (Melo et al., 2013), with about 700,000 ha forest restored between 2011 and 2015 (Crouzeilles et al., 2019). Yet, Bicudo da Silva et al. (2023) showed that between 2001-2015 there was a process called “forest transition” (declines in forest cover cease and recoveries in forest cover begin) (Rudel et al., 2005), due to the stagnation of agricultural activities, the emergence of non-agricultural rural activities, and the decrease in precipitation leading to soil abandonment and favoring regeneration.

In Argentina, the percentage of forest has been reduced linearly since the 1990s, with the combined effect of the advance of small-scale agriculture associated with population growth and road construction in some areas, and the increase of monospecific forest plantations incentivized by government subsidies and the participation of large timber companies (Izquierdo et al., 2008). The forest loss rate was lower during 2005-2015, potentially because of the effect of the certified wood market in this region and the approval of the National Forest Law and the implementation of the National Fund for the Enrichment and Conservation of Native Forests (FVSA & WWF, 2017). However, forest loss increased in the last period (2015-2020) most likely due to higher levels of economic growth and the impact of long-term police on the expansion of agriculture and cattle raising in this province (Mohebalian et al., 2022). Paraguay showed the highest rates of deforestation of the entire Atlantic Forest between 1986-2005 due to the massive expansion of agriculture. However, since the creation of the Zero Deforestation Law and the implementation of associated mechanisms, there has been a recent stabilization of vegetation loss (Da Ponte et al., 2017; FVSA & WWF, 2017).

### 4.4 Core and edge area

Our results showed that 50% of the remaining vegetation is under the effect of a 90 m edge, and about 75% is under the effect of a 240 m edge and almost 90% is under the effect of a 500 m edge, results very similar to Haddad et al. (2015). Over time, there was an increase in vegetation located less than 90 m from the edges, revealing a pronounced edge effect threshold in the AF. Below this threshold, there is an intensification of edge effects, and above it, there is a decrease in the amount of vegetation core. This threshold is probably associated with the massive number and small average size of fragments we detected. Importantly, small fragments are more subject to edge effects due to their size and shape (Fahrig, 2003). The edge effect changes the AF landscape features such as microclimate and carbon cycle (Magnago et al., 2017, 2015) depending on the fragment shape (Banks-Leite et al., 2013) and the matrix effect (Adorno et al., 2021). In that regard, numerous studies have demonstrated the negative effects of edge changes for epiphyte plants, small mammalian, and birds in the AF (de la Sancha et al., 2023; Morante-Filho et al., 2018; Parra-Sanchez and Banks-Leite, 2020). Added to that, Pivello et al. (2021) identified that AF is highly fire-sensitive, which changes the conditions of the edges and vice versa. Some measures such as forested or agroforestry matrices and strips of trees being planted, forming a buffer around the remaining fragments can reduce the edge effect (Gama-Rodrigues et al., 2021; Tavares et al., 2019).

### 4.5 Functional connectivity and mean isolation

Functional connectivity and isolation had similar response patterns over time, with their worst values between 1990 and 2000, but from 2005 onwards there were clear signs of improvement. The vegetation amount has not changed noticeably since 2005, and this improvement was due to the appearance of new fragments that increased the connectivity of the landscape, probably through stepping stones. In this way, small fragments (<50 ha, which represents 97% of AF remnants) play a fundamental role in keeping large fragments connected, even more important for species that can cross the matrix (Diniz et al., 2021). Furthermore, NV plays a key role in decreasing the isolation of the remnants, although there may be fewer forest-specialist species that use this type of vegetation, it can be critical to maintaining AF connectivity (Lyra-Jorge et al., 2010).

However, practices such as agroecology and forestry can increase the connectivity by increasing the permeability of the matrix (Tubenchlak et al., 2021). In addition, The Atlantic Forest Restoration Pact and Rural Environment Registry (CAR) police are a great opportunity to create and improve ecological corridors (da Silva et al., 2023; Melo et al., 2013). Finally, although connectivity and isolation were not apparently sensitive to the roads and railways effect, this lack of sensitivity may be due to short-distance divisions into FV or NV fragments, as the additional cost that these linear structures cause, preventing animals from crossing short distances (Martinez Pardo et al., 2023), were not considered. Thus, it is essential to propose fauna passages for improving landscape permeability to maintain wildlife gene flow and reduce roadkill (Cassimiro et al., 2023; Teixeira et al., 2022; Zimmermann Teixeira et al., 2017).

### 4.6 Protected areas and indigenous territories

Alarmingly, our results showed that the proportion of PA (10% for FV and 8.3% for NV) is far below the targets (30% land surface by 2030) of the post-2020 Global Biodiversity Framework (Jung et al., 2021). Moreover, these values are higher but consistent with those found in previous years, such as 9.3% by Ribeiro et al. (2009) and 9% by Rezende et al. (2018). We highlight that IT, despite not being PA, has proven to be fundamental for forest restoration in AF (Benzeev et al., 2023). Noticeably, 70% and 90% of vegetation is more than 10 km distant from PA and IT. Our findings are more alarming than those found by Ribeiro et al. (2009). Forest formation has the largest area in PA (8.2%) and smaller for IT (1.5%). This result is expected because of its large contribution to AF composition (62.1%) because forests have been commonly the main target for PA creation. In addition, *restinga* and mangroves had a high overlap with PA and TI (40%), due to the high density of these protective measures on the Brazilian coast, especially in Serra do Mar. However, despite this high proportion of protection, these ecosystems have faced many threats in recent decades, which can affect several functions of ecosystems and local populations (Diniz et al., 2019). Savanna formation was critical to ensuring connectivity, however, this class has the lowest proportion of PA and TI (4.7%) despite representing 23% of the amount of vegetation, possibly because this vegetation formation is not guaranteed by specific protection laws. Since deforestation outside PA and IT has been lower than in private rural areas (da Silva et al., 2023), these areas are essential to ensure biodiversity conservation (Avigliano et al., 2019; Benzeev et al., 2023). Therefore, it is necessary to create new PA and IT, and strengthen the connection network between existing ones, as well as restrictions in their surroundings to promote the restoration of vegetation.

## 5. Conclusion

To our knowledge, this is the first work that analyzed the spatiotemporal dynamics of the entire AF landscape structure through multiple landscape metrics, considering a broad tri-national delimitation, only forest vegetation and both forest and other natural vegetation, and the effect of roads and railways. Our findings allow a detailed understanding of the habitat fragmentation process in the AF in the last three and half decades. The number of FV fragments has increased, which comes accompanied by an important increment of vegetation. Besides that, NV—fundamental to promote connectivity—is far from being under enough protection. Overall, the fragmentation scenarios in Argentina, Brazil, and Paraguay are equally worrying (97% of fragments are very small and 60% are under edge effect). We also highlight the substantial effect of roads and railways on breaking large FV fragments apart, likely disrupting the functional connectivity of several ecological processes. These results lead us to reinforce the need for conservation and restoration actions, such as investing in implementing conservation plans for large fragments, promoting the connectivity of small fragments, managing the matrix to minimize edge effects and improve connectivity, and leading restoration actions in key areas, such as large and isolated fragments and indigenous territory. Added to this, we highlight the importance of planning and building fauna passages to improve landscape connectivity and reduce wildlife roadkill. Finally, the protection legislation implemented in mid-2005, combined with the restoration initiatives started in 2009, and the implementation of the CAR in 2012 appear to be having an effect in starting a process of AF restoration. The continuity and expansion of these measures are essential to guarantee the continuity of this AF process in the future, given the new threats of climate change and the expansion of urban and agricultural areas.

## Funding information

MHV was supported by grants #2013/02883-7, #2017/09676-8, and #2022/01899-6, São Paulo Research Foundation (FAPESP), and CAPES (grant numbers 88887.513979/2020-00 and 1588183). RLM was supported by Bryce Carmine and Anne Carmine (née Percival), through the Massey University Foundation. CHG is a research fellow of the National Council for Scientific and Technological Development - CNPq (grant 311209/2021-1). JEFO was supported by FAPESP projects #2014/23132-2, and #2021/02132-2. MCR was supported by FAPESP (processes #2013/50421-2; #2020/01779-5; #2021/08322-3; #2021/08534-0; #2021/10195-0; #2021/10639-5; #2022/10760-1) and CNPq (processes #442147/2020-1; #440145/2022-8; #402765/2021-4; #313016/2021-6; #440145/2022-8), and São Paulo State University - UNESP for their financial support. This study was financed in part by CAPES Brazil – Finance Code 001. This study is also part of the *Center for Research on Biodiversity Dynamics and Climate Change*, which is financed by FAPESP.

## CRediT authorship contribution statement

**MHV**: Conceptualization, Data curation, Formal analysis, Investigation, Methodology, Software, Visualization, Writing— original draft, Writing—review and editing. **RLM**: Conceptualization, Writing—review and editing. **BBN**: Conceptualization, Methodology, Software, Writing—review and editing. **JEFO**: Conceptualization, Writing—review and editing. **VT**: Conceptualization, Writing—review and editing. **RB**: Writing—review and editing. **CDA**: Writing—review and editing. **MRR**: Writing—review and editing. **CHG**: Methodology, Software, Writing—review and editing. **MCR**: Conceptualization, Funding acquisition, Investigation, Methodology, Project administration, Resources, Software, Supervision, Writing—review and editing. All authors gave final approval for publication and agreed to be held accountable for the work performed therein.

## Declaration of competing interest

The authors declare that they have no known competing financial interests or personal relationships that could have appeared to influence the work reported in this paper.

## Data availability

Code provided in GitHub (https://github.com/LEEClab/ms-atlantic-forest-spatiotemporal-dynamics). Data and code are provided in Open Science Files (OSF) (https://doi.org/10.17605/OSF.IO/RFWBZ).

## Acknowledgements

We thank Luis Fernando Guedes Pinto from SOS Mata Atlântica for assistance with MapBiomas data and Atlantic Forest delimitation. We also thank Alexandre Uezu for valuable contributions to the manuscript. We thank all members from Spatial Ecology and Conservation Lab (LEEC) for their help with fruitful discussions and suggestions, especially Rafael Souza. MHV thanks João Giovanelli, Rodrigo Nobre, Rafael Fávero, and Thiago Gonçalves-Souza for their valuable proposals for data analysis and discussion of results, Alejandro R. Vargas by vegetation cover in Argentina and Paraguay, and Maurício E. Graipel by for pointing out the role of legislation in large-scale vegetation dynamics.

## Notes

https://github.com/LEEClab/ms-atlantic-forest-spatiotemporal-dynamics/tree/main

### Competing Interest Statement

The authors have declared no competing interest.

### Summary of Updates

- Correction of the NV percentage value in the summary and text - Correction of the citation of the number of plants in the Atlantic Forest "18,000 species of plants (Flora e Funga do Brasil, 2023)" - Correction of natural vegetation cover in the discussion - Correction of references

