## supplementary_material for "The Atlantic Forest of South America: spatiotemporal dynamics of remaining vegetation and implications for conservation"

#### Data

Roads and railways data were downloaded from official geospatial databases from three countries: Brazil (Instituto Brasileiro de Geografia e Estatística – IBGE; IBGE, 2021), Argentina (Instituto Geográfico Nacional – IGN; IGN, 2022), and Paraguay (Instituto Nacional de Estadística – INE; INE, 2022). The data comprised 14,072 km of railways and 124,187 km of roads, with a total of 138,259 km (Fig. S11). We did not find official railway data for Paraguay, so there may be an underestimation of this effect for this country. We couldn't differentiate the roads and railways by year, as it's almost impossible to find that information, so we used the same layer to trim all the fragments in all years. We assume that this may have overestimated the effect of roads in the past when some of these roads or railways did not exist or were not paved yet. For Brazil, we selected paved, operational, and constructed roads, and railways that were selected by relative surface position, and train section. The road and railways layers were rasterized using a parameter that creates densified lines, i.e., all cells touched by the line were included as data for rasterization, which resulted in more densified lines. This guaranteed that the roads and railways would take up space and trimmed the fragments. After rasterizing the lines, the raster covered 524,147 ha (0.32% from AF delimitation). Thus, we trimmed the fragments of vegetation from the rasterized data.

The Protected Areas (PA) were downloaded from Protected Planet (UNEP-WCMC and IUCN, 2022) for the IUCN categories of protected areas (“Ia”, “Ib”, “II”, “III”, and “IV”) following (Rezende et al., 2018), which comprises 986 reserves (4,620,245 ha; 2.84% from AF delimitation) (Fig. S12a). These IUCN categories encompass multiply protection categories for Argentina (Municipal Nature Park, National Park, Nature Monument, Private Refuge, Private Wildlife Refuge, Provincial Park, Strict Nature Reserve, Wilderness Nature Reserve, and Wildlife Reserve), Brazil (Area of Relevant Ecological Interest, Biological Reserve, Ecological Station, Natural Heritage Private Reserve, Natural Monument, Park, Ramsar Site, Wetland of International Importance, Wildlife Refuge), and Paraguay (National Park, Natural Private Reserve, Natural Reserve, Scientific Monument, and Scientific Reserve). The Indigenous Territories (IT) were downloaded for Brazil (Fundação Nacional dos Povos Indígenas; FUNAI, 2020) and Paraguay (Tierras Indígenas, 2022), which comprises 1023 territories (1,324,973 ha; 0.81% from AF delimitation) (Fig. S12b). We did not find official IT data for Argentina; therefore, we cannot analyze the contribution of IT to this country.

### Figures

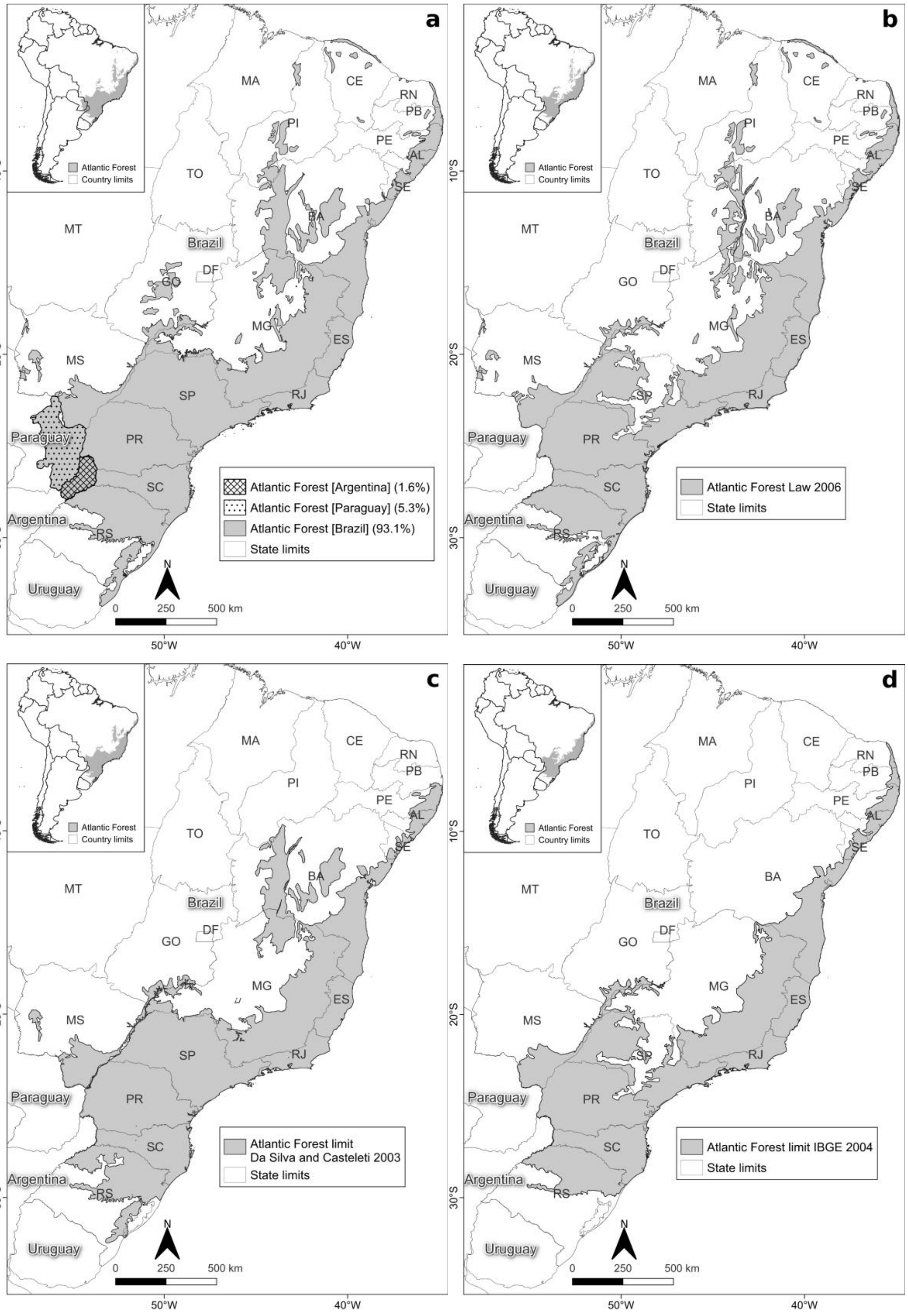

a
Atlantic Forest
Country limits
MA
CE
RN
PB
PI
PE
AL
TO
SE
BA
MT
Brazil
DF
GO
MG
ES
MS
SP
RJ
Paraguay
PR
SC
Argentina
RS
Uruguay
Atlantic Forest [Argentina] (1.6%)
Atlantic Forest [Paraguay] (5.3%)
Atlantic Forest [Brazil] (93.1%)
State limits
N
0
250
500 km
10°S
20°S
30°S
50°W
40°W
b
Atlantic Forest Law 2006
State limits
c
Atlantic Forest limit
Da Silva and Casteleti 2003
State limits
d
Atlantic Forest limit IBGE 2004
State limits

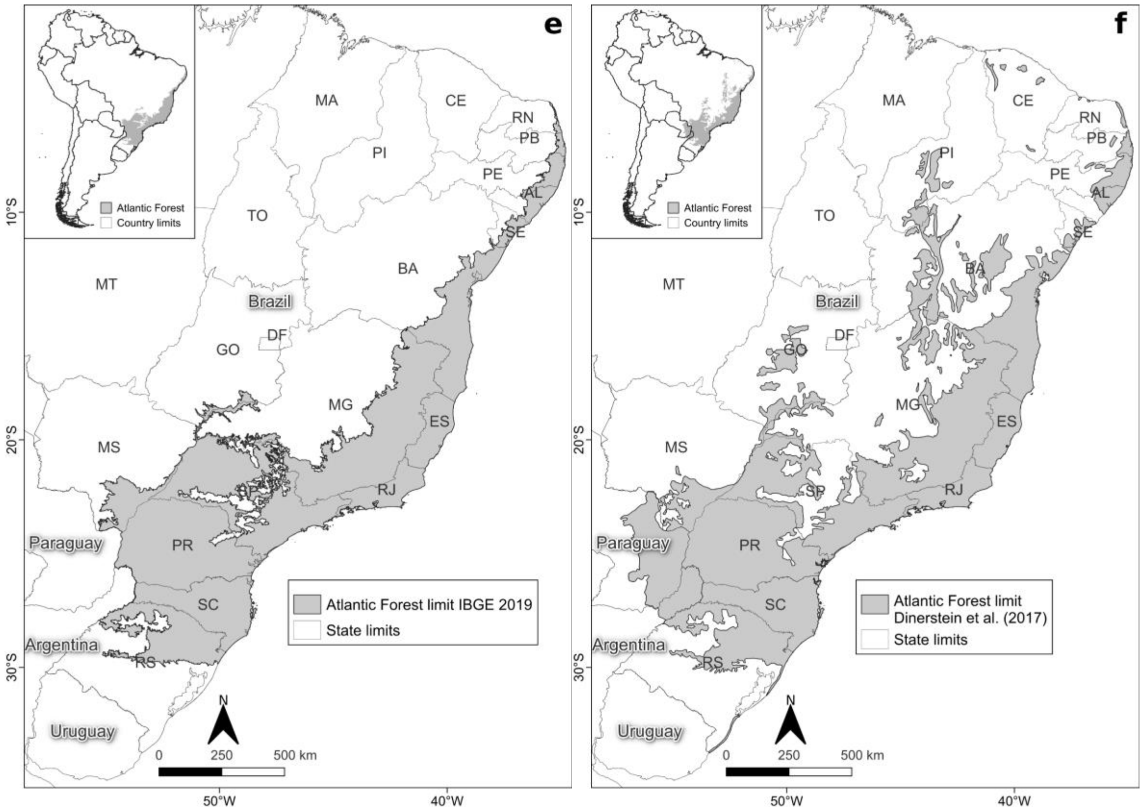


**Figure S1**. Atlantic Forest delimitation, adapted from Muylaert et al. (2018) (a). Atlantic Forest delimitations used to create the “integrative delimitation” incorporating different types of ecosystems: (b) delimitation defined by Brazilian legislation (Federal Decree No. 750/93 and Atlantic Forest law No. 11 428, of December 22, 2006); (c) delimitation defined by Da Silva and Casteleti (2003); (d) AF delimitation defined by IBGE (2004), (e) AF delimitation defined by IBGE (2019) and; (f) AF delimitation defined by (Dinerstein et al., 2017) and used in the Ecoregions 2017®.

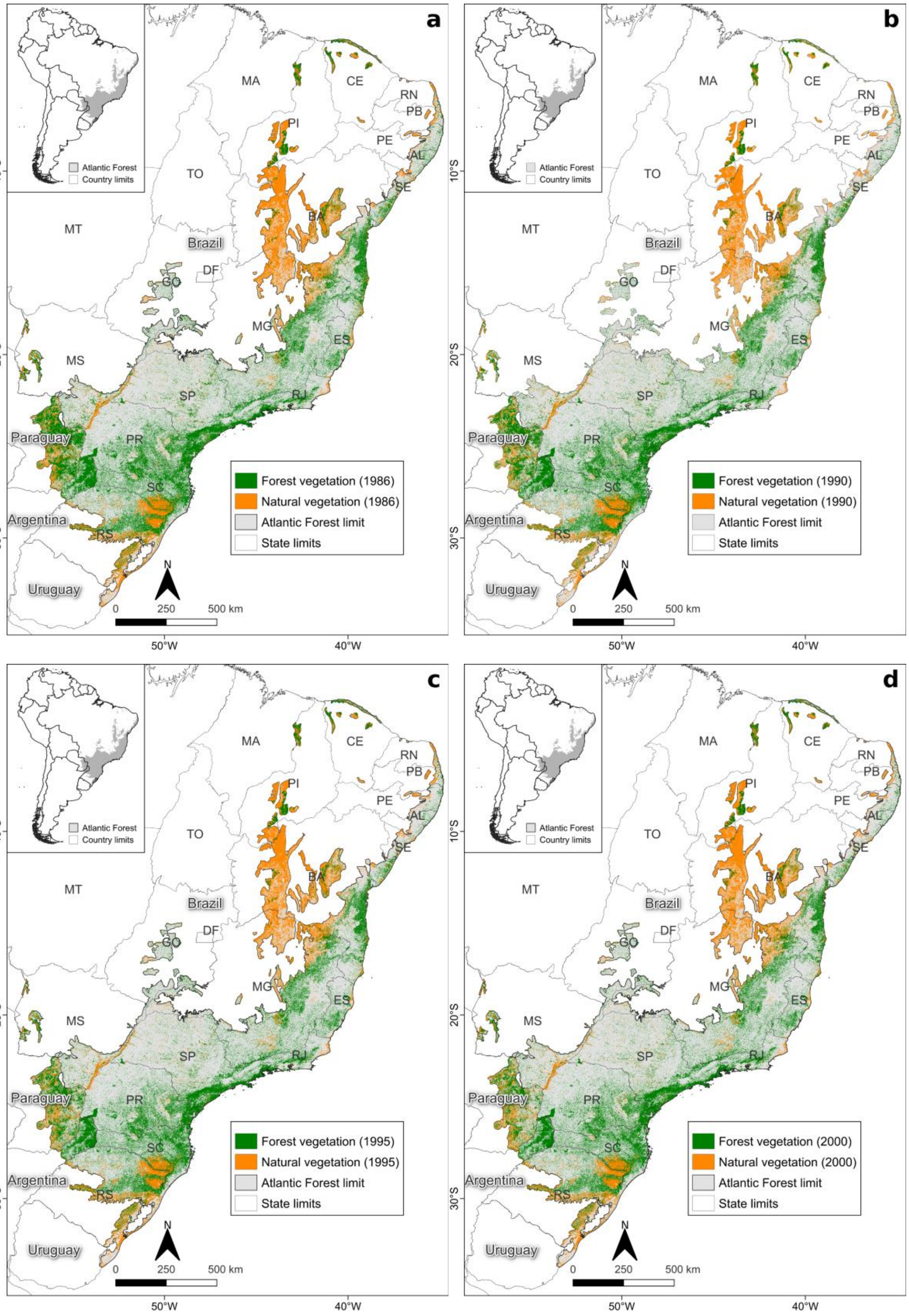
a
b
c
d
Atlantic Forest
Country limits
MA
CE
RN
PB
PI
PE
AL
TO
SE
BA
MT
Brazil
DF
GO
MG
ES
MS
SP
RJ
Paraguay
PR
SC
Argentina
RS
Uruguay
N
0
250
500 km
10°S
20°S
30°S
50°W
40°W
Forest vegetation (1986)
Natural vegetation (1986)
Forest vegetation (1990)
Natural vegetation (1990)
Forest vegetation (1995)
Natural vegetation (1995)
Forest vegetation (2000)
Natural vegetation (2000)
Atlantic Forest limit
State limits

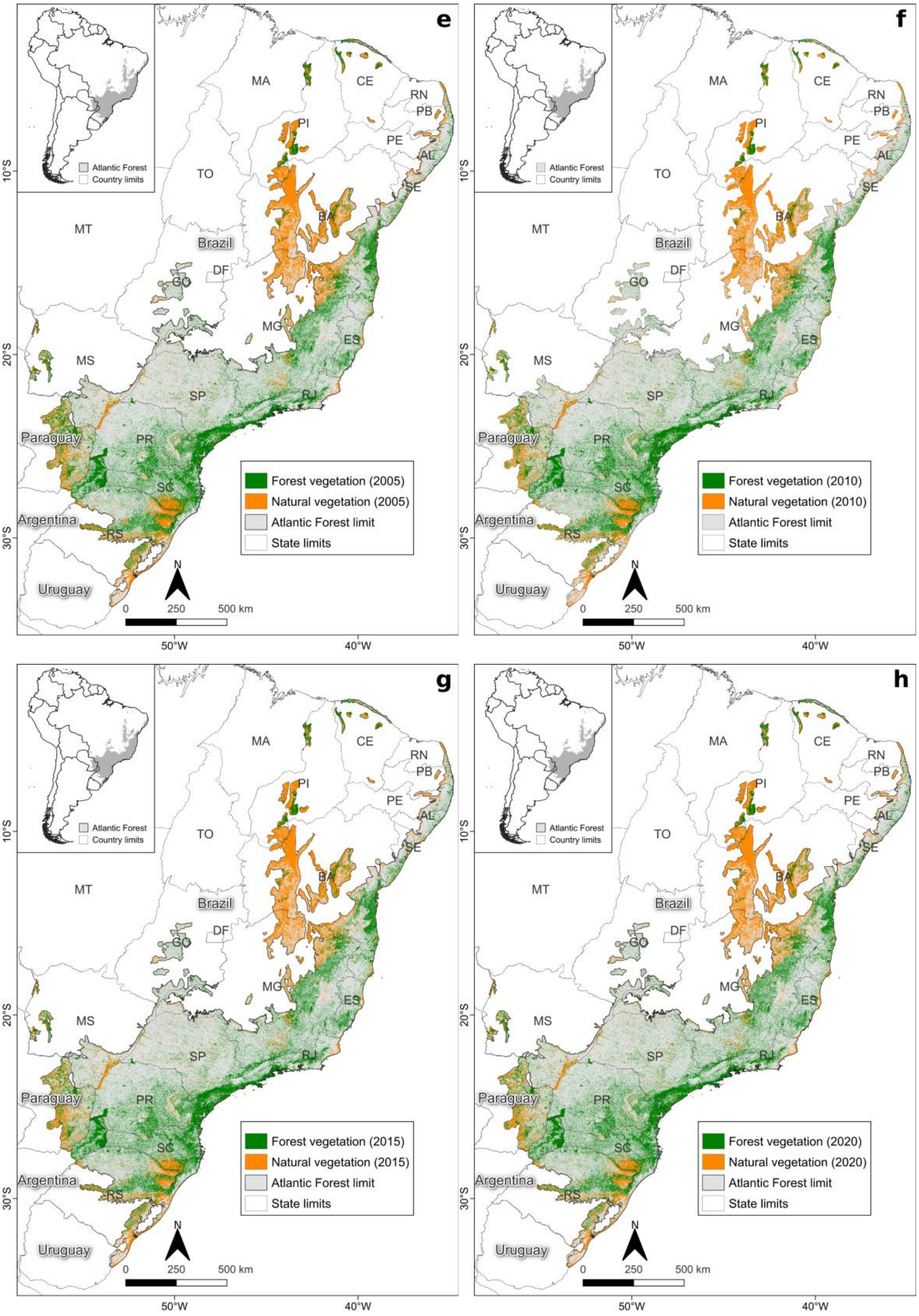


**Figure S2**. FV and NV cover for the entire Atlantic Forest from 1986-2020 (a-h).

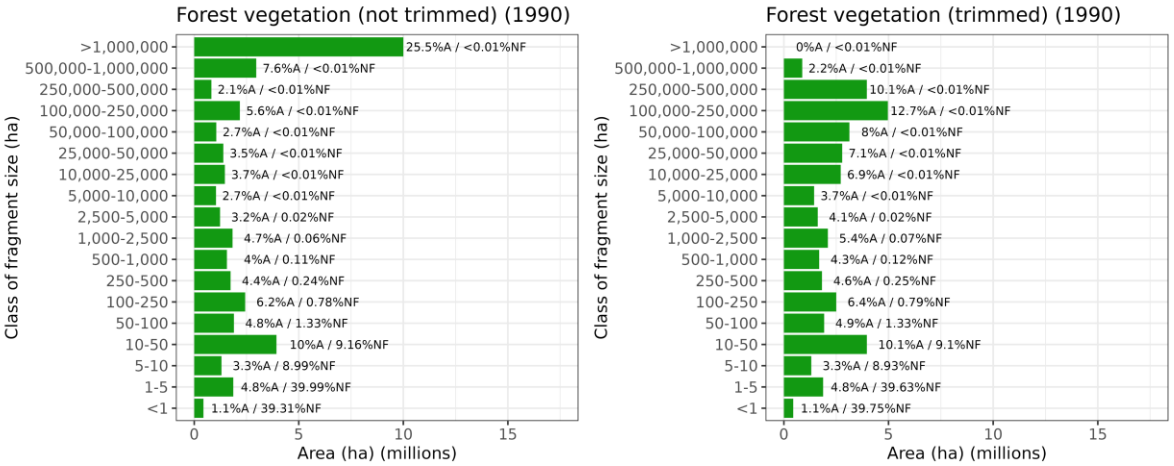
Forest vegetation (not trimmed) (1990)
Forest vegetation (trimmed) (1990)
Class of fragment size (ha)
>1,000,000
500,000-1,000,000
250,000-500,000
100,000-250,000
50,000-100,000
25,000-50,000
10,000-25,000
5,000-10,000
2,500-5,000
1,000-2,500
500-1,000
250-500
100-250
50-100
10-50
5-10
1-5
<1
25.5%A / <0.01%NF
7.6%A / <0.01%NF
2.1%A / <0.01%NF
5.6%A / <0.01%NF
2.7%A / <0.01%NF
3.5%A / <0.01%NF
3.7%A / <0.01%NF
2.7%A / <0.01%NF
3.2%A / 0.02%NF
4.7%A / 0.06%NF
4%A / 0.11%NF
4.4%A / 0.24%NF
6.2%A / 0.78%NF
4.8%A / 1.33%NF
10%A / 9.16%NF
3.3%A / 8.99%NF
4.8%A / 39.99%NF
1.1%A / 39.31%NF
0%A / <0.01%NF
2.2%A / <0.01%NF
10.1%A / <0.01%NF
12.7%A / <0.01%NF
8%A / <0.01%NF
7.1%A / <0.01%NF
6.9%A / <0.01%NF
3.7%A / <0.01%NF
4.1%A / 0.02%NF
5.4%A / 0.07%NF
4.3%A / 0.12%NF
4.6%A / 0.25%NF
6.4%A / 0.79%NF
4.9%A / 1.33%NF
10.1%A / 9.1%NF
3.3%A / 8.93%NF
4.8%A / 39.63%NF
1.1%A / 39.75%NF
0 5 10 15
Area (ha) (millions)

96

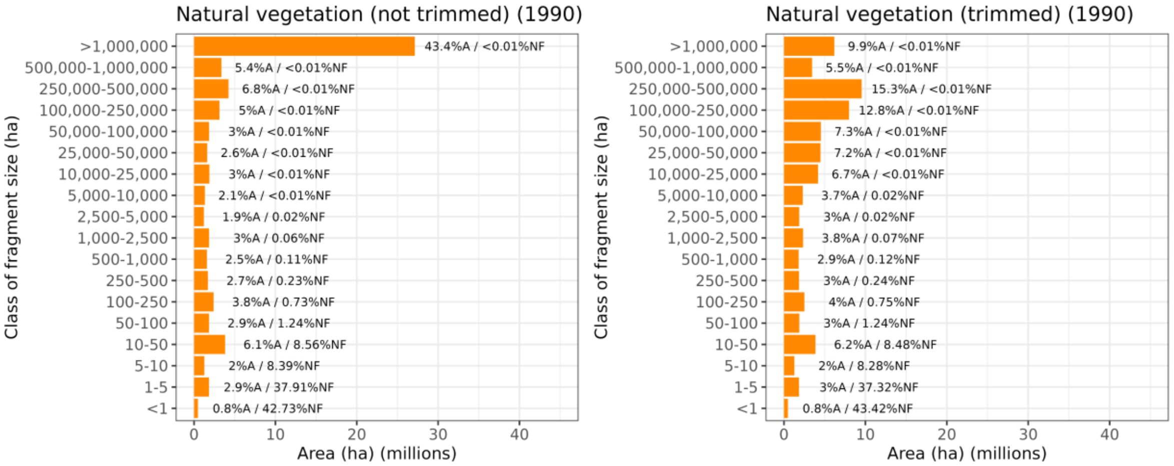
Natural vegetation (not trimmed) (1990)
Natural vegetation (trimmed) (1990)
Class of fragment size (ha)
>1,000,000
500,000-1,000,000
250,000-500,000
100,000-250,000
50,000-100,000
25,000-50,000
10,000-25,000
5,000-10,000
2,500-5,000
1,000-2,500
500-1,000
250-500
100-250
50-100
10-50
5-10
1-5
<1
43.4%A / <0.01%NF
5.4%A / <0.01%NF
6.8%A / <0.01%NF
5%A / <0.01%NF
3%A / <0.01%NF
2.6%A / <0.01%NF
3%A / <0.01%NF
2.1%A / <0.01%NF
1.9%A / 0.02%NF
3%A / 0.06%NF
2.5%A / 0.11%NF
2.7%A / 0.23%NF
3.8%A / 0.73%NF
2.9%A / 1.24%NF
6.1%A / 8.56%NF
2%A / 8.39%NF
2.9%A / 37.91%NF
0.8%A / 42.73%NF
9.9%A / <0.01%NF
5.5%A / <0.01%NF
15.3%A / <0.01%NF
12.8%A / <0.01%NF
7.3%A / <0.01%NF
7.2%A / <0.01%NF
6.7%A / <0.01%NF
3.7%A / 0.02%NF
3%A / 0.02%NF
3.8%A / 0.07%NF
2.9%A / 0.12%NF
3%A / 0.24%NF
4%A / 0.75%NF
3%A / 1.24%NF
6.2%A / 8.48%NF
2%A / 8.28%NF
3%A / 37.32%NF
0.8%A / 43.42%NF
0 10 20 30 40
Area (ha) (millions)

97

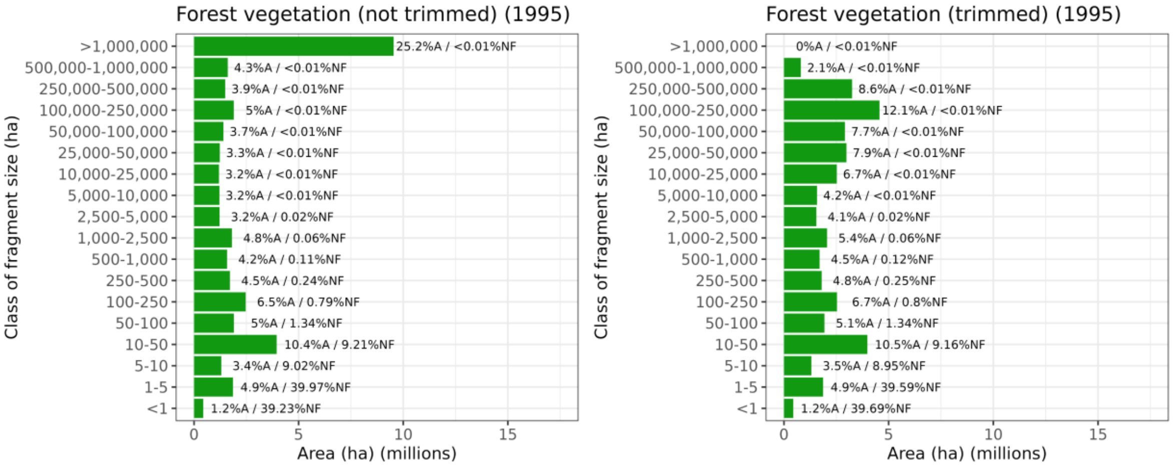
Forest vegetation (not trimmed) (1995)
Forest vegetation (trimmed) (1995)
Class of fragment size (ha)
>1,000,000
500,000-1,000,000
250,000-500,000
100,000-250,000
50,000-100,000
25,000-50,000
10,000-25,000
5,000-10,000
2,500-5,000
1,000-2,500
500-1,000
250-500
100-250
50-100
10-50
5-10
1-5
<1
25.2%A / <0.01%NF
4.3%A / <0.01%NF
3.9%A / <0.01%NF
5%A / <0.01%NF
3.7%A / <0.01%NF
3.3%A / <0.01%NF
3.2%A / <0.01%NF
3.2%A / <0.01%NF
3.2%A / 0.02%NF
4.8%A / 0.06%NF
4.2%A / 0.11%NF
4.5%A / 0.24%NF
6.5%A / 0.79%NF
5%A / 1.34%NF
10.4%A / 9.21%NF
3.4%A / 9.02%NF
4.9%A / 39.97%NF
1.2%A / 39.23%NF
0%A / <0.01%NF
2.1%A / <0.01%NF
8.6%A / <0.01%NF
12.1%A / <0.01%NF
7.7%A / <0.01%NF
7.9%A / <0.01%NF
6.7%A / <0.01%NF
4.2%A / <0.01%NF
4.1%A / 0.02%NF
5.4%A / 0.06%NF
4.5%A / 0.12%NF
4.8%A / 0.25%NF
6.7%A / 0.8%NF
5.1%A / 1.34%NF
10.5%A / 9.16%NF
3.5%A / 8.95%NF
4.9%A / 39.59%NF
1.2%A / 39.69%NF
0 5 10 15
Area (ha) (millions)

98

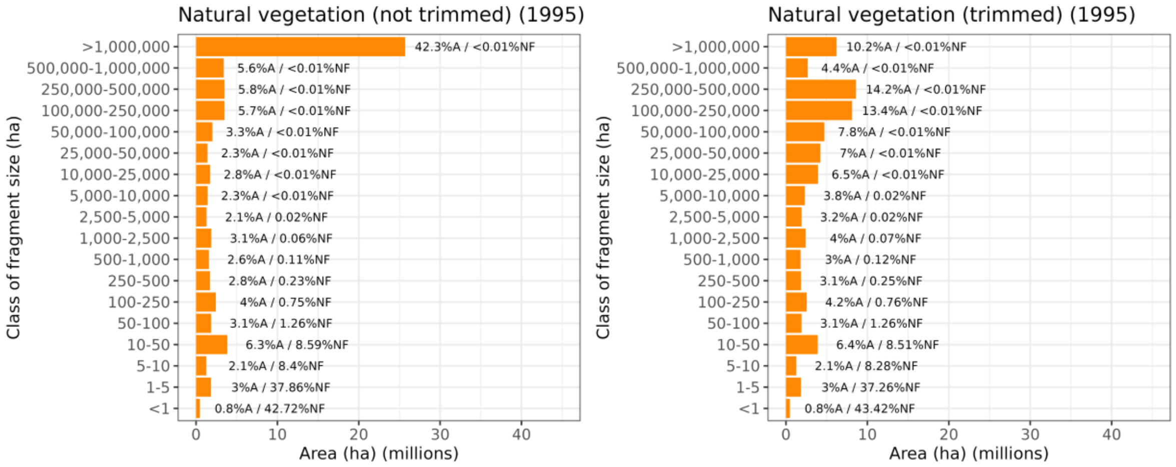
Natural vegetation (not trimmed) (1995)
Class of fragment size (ha)
>1,000,000
500,000-1,000,000
250,000-500,000
100,000-250,000
50,000-100,000
25,000-50,000
10,000-25,000
5,000-10,000
2,500-5,000
1,000-2,500
500-1,000
250-500
100-250
50-100
10-50
5-10
1-5
<1
42.3%A / <0.01%NF
5.6%A / <0.01%NF
5.8%A / <0.01%NF
5.7%A / <0.01%NF
3.3%A / <0.01%NF
2.3%A / <0.01%NF
2.8%A / <0.01%NF
2.3%A / <0.01%NF
2.1%A / 0.02%NF
3.1%A / 0.06%NF
2.6%A / 0.11%NF
2.8%A / 0.23%NF
4%A / 0.75%NF
3.1%A / 1.26%NF
6.3%A / 8.59%NF
2.1%A / 8.4%NF
3%A / 37.86%NF
0.8%A / 42.72%NF
0 10 20 30 40
Area (ha) (millions)
Natural vegetation (trimmed) (1995)
Class of fragment size (ha)
10.2%A / <0.01%NF
4.4%A / <0.01%NF
14.2%A / <0.01%NF
13.4%A / <0.01%NF
7.8%A / <0.01%NF
7%A / <0.01%NF
6.5%A / <0.01%NF
3.8%A / 0.02%NF
3.2%A / 0.02%NF
4%A / 0.07%NF
3%A / 0.12%NF
3.1%A / 0.25%NF
4.2%A / 0.76%NF
3.1%A / 1.26%NF
6.4%A / 8.51%NF
2.1%A / 8.28%NF
3%A / 37.26%NF
0.8%A / 43.42%NF
0 10 20 30 40
Area (ha) (millions)

99

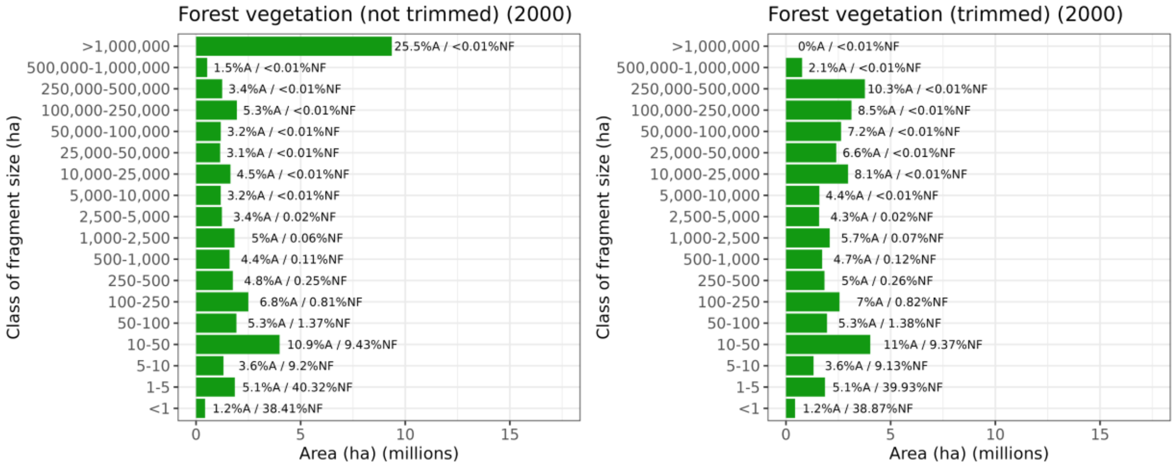
Forest vegetation (not trimmed) (2000)
Class of fragment size (ha)
>1,000,000
500,000-1,000,000
250,000-500,000
100,000-250,000
50,000-100,000
25,000-50,000
10,000-25,000
5,000-10,000
2,500-5,000
1,000-2,500
500-1,000
250-500
100-250
50-100
10-50
5-10
1-5
<1
25.5%A / <0.01%NF
1.5%A / <0.01%NF
3.4%A / <0.01%NF
5.3%A / <0.01%NF
3.2%A / <0.01%NF
3.1%A / <0.01%NF
4.5%A / <0.01%NF
3.2%A / <0.01%NF
3.4%A / 0.02%NF
5%A / 0.06%NF
4.4%A / 0.11%NF
4.8%A / 0.25%NF
6.8%A / 0.81%NF
5.3%A / 1.37%NF
10.9%A / 9.43%NF
3.6%A / 9.2%NF
5.1%A / 40.32%NF
1.2%A / 38.41%NF
0 5 10 15
Area (ha) (millions)
Forest vegetation (trimmed) (2000)
Class of fragment size (ha)
0%A / <0.01%NF
2.1%A / <0.01%NF
10.3%A / <0.01%NF
8.5%A / <0.01%NF
7.2%A / <0.01%NF
6.6%A / <0.01%NF
8.1%A / <0.01%NF
4.4%A / <0.01%NF
4.3%A / 0.02%NF
5.7%A / 0.07%NF
4.7%A / 0.12%NF
5%A / 0.26%NF
7%A / 0.82%NF
5.3%A / 1.38%NF
11%A / 9.37%NF
3.6%A / 9.13%NF
5.1%A / 39.93%NF
1.2%A / 38.87%NF
0 5 10 15
Area (ha) (millions)

100

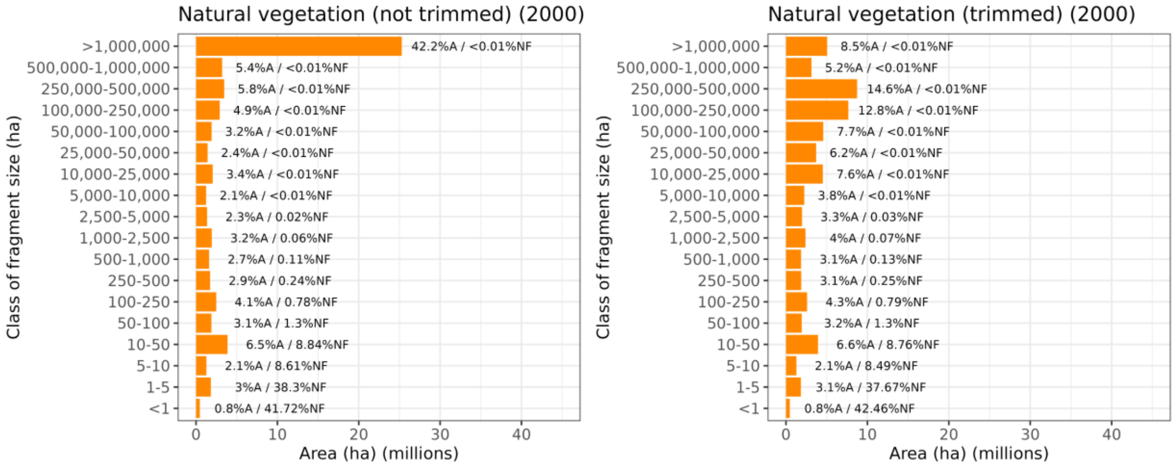
Natural vegetation (not trimmed) (2000)
Class of fragment size (ha)
>1,000,000
500,000-1,000,000
250,000-500,000
100,000-250,000
50,000-100,000
25,000-50,000
10,000-25,000
5,000-10,000
2,500-5,000
1,000-2,500
500-1,000
250-500
100-250
50-100
10-50
5-10
1-5
<1
42.2%A / <0.01%NF
5.4%A / <0.01%NF
5.8%A / <0.01%NF
4.9%A / <0.01%NF
3.2%A / <0.01%NF
2.4%A / <0.01%NF
3.4%A / <0.01%NF
2.1%A / <0.01%NF
2.3%A / 0.02%NF
3.2%A / 0.06%NF
2.7%A / 0.11%NF
2.9%A / 0.24%NF
4.1%A / 0.78%NF
3.1%A / 1.3%NF
6.5%A / 8.84%NF
2.1%A / 8.61%NF
3%A / 38.3%NF
0.8%A / 41.72%NF
0 10 20 30 40
Area (ha) (millions)
Natural vegetation (trimmed) (2000)
Class of fragment size (ha)
8.5%A / <0.01%NF
5.2%A / <0.01%NF
14.6%A / <0.01%NF
12.8%A / <0.01%NF
7.7%A / <0.01%NF
6.2%A / <0.01%NF
7.6%A / <0.01%NF
3.8%A / <0.01%NF
3.3%A / 0.03%NF
4%A / 0.07%NF
3.1%A / 0.13%NF
3.1%A / 0.25%NF
4.3%A / 0.79%NF
3.2%A / 1.3%NF
6.6%A / 8.76%NF
2.1%A / 8.49%NF
3.1%A / 37.67%NF
0.8%A / 42.46%NF
0 10 20 30 40
Area (ha) (millions)

101

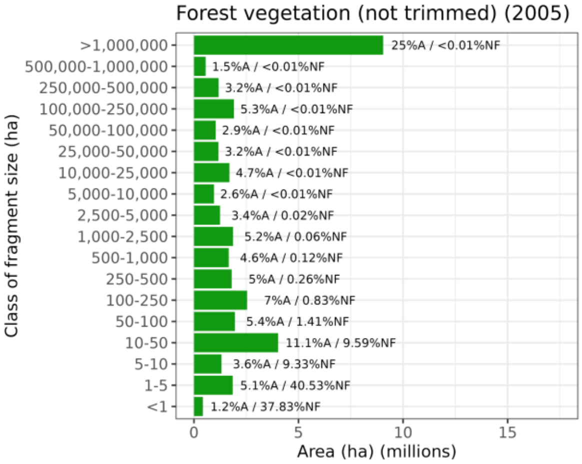
Forest vegetation (not trimmed) (2005)
Class of fragment size (ha)
Area (ha) (millions)
>1,000,000
500,000-1,000,000
250,000-500,000
100,000-250,000
50,000-100,000
25,000-50,000
10,000-25,000
5,000-10,000
2,500-5,000
1,000-2,500
500-1,000
250-500
100-250
50-100
10-50
5-10
1-5
<1
25%A / <0.01%NF
1.5%A / <0.01%NF
3.2%A / <0.01%NF
5.3%A / <0.01%NF
2.9%A / <0.01%NF
3.2%A / <0.01%NF
4.7%A / <0.01%NF
2.6%A / <0.01%NF
3.4%A / 0.02%NF
5.2%A / 0.06%NF
4.6%A / 0.12%NF
5%A / 0.26%NF
7%A / 0.83%NF
5.4%A / 1.41%NF
11.1%A / 9.59%NF
3.6%A / 9.33%NF
5.1%A / 40.53%NF
1.2%A / 37.83%NF
0
5
10
15

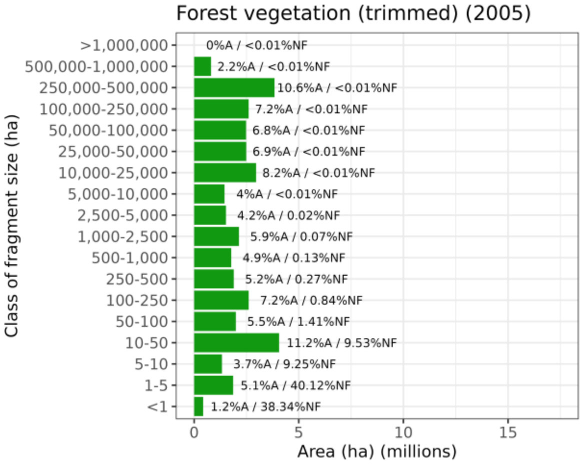
Forest vegetation (trimmed) (2005)
Class of fragment size (ha)
Area (ha) (millions)
>1,000,000
500,000-1,000,000
250,000-500,000
100,000-250,000
50,000-100,000
25,000-50,000
10,000-25,000
5,000-10,000
2,500-5,000
1,000-2,500
500-1,000
250-500
100-250
50-100
10-50
5-10
1-5
<1
0%A / <0.01%NF
2.2%A / <0.01%NF
10.6%A / <0.01%NF
7.2%A / <0.01%NF
6.8%A / <0.01%NF
6.9%A / <0.01%NF
8.2%A / <0.01%NF
4%A / <0.01%NF
4.2%A / 0.02%NF
5.9%A / 0.07%NF
4.9%A / 0.13%NF
5.2%A / 0.27%NF
7.2%A / 0.84%NF
5.5%A / 1.41%NF
11.2%A / 9.53%NF
3.7%A / 9.25%NF
5.1%A / 40.12%NF
1.2%A / 38.34%NF
0
5
10
15

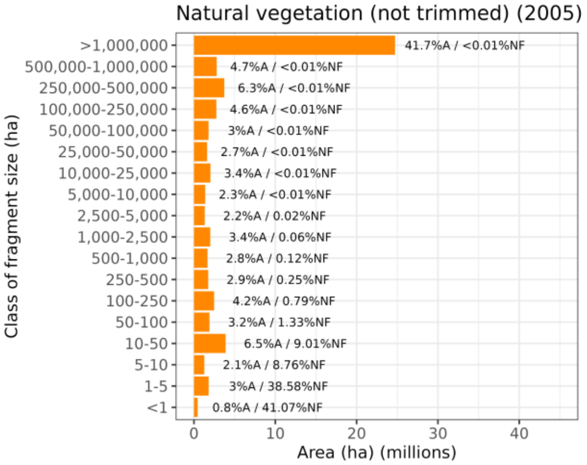
Natural vegetation (not trimmed) (2005)
Class of fragment size (ha)
Area (ha) (millions)
>1,000,000
500,000-1,000,000
250,000-500,000
100,000-250,000
50,000-100,000
25,000-50,000
10,000-25,000
5,000-10,000
2,500-5,000
1,000-2,500
500-1,000
250-500
100-250
50-100
10-50
5-10
1-5
<1
41.7%A / <0.01%NF
4.7%A / <0.01%NF
6.3%A / <0.01%NF
4.6%A / <0.01%NF
3%A / <0.01%NF
2.7%A / <0.01%NF
3.4%A / <0.01%NF
2.3%A / <0.01%NF
2.2%A / 0.02%NF
3.4%A / 0.06%NF
2.8%A / 0.12%NF
2.9%A / 0.25%NF
4.2%A / 0.79%NF
3.2%A / 1.33%NF
6.5%A / 9.01%NF
2.1%A / 8.76%NF
3%A / 38.58%NF
0.8%A / 41.07%NF
0
10
20
30
40

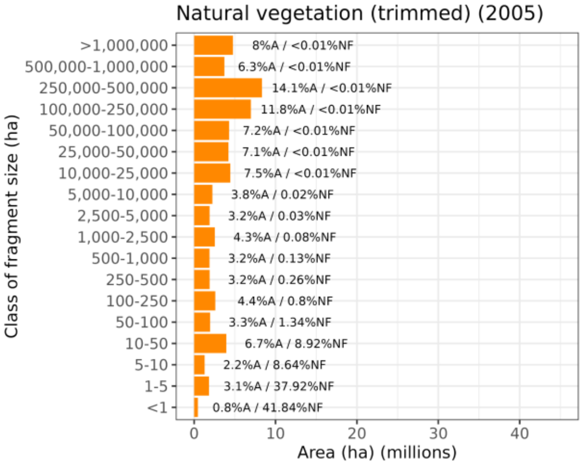
Natural vegetation (trimmed) (2005)
Class of fragment size (ha)
Area (ha) (millions)
>1,000,000
500,000-1,000,000
250,000-500,000
100,000-250,000
50,000-100,000
25,000-50,000
10,000-25,000
5,000-10,000
2,500-5,000
1,000-2,500
500-1,000
250-500
100-250
50-100
10-50
5-10
1-5
<1
8%A / <0.01%NF
6.3%A / <0.01%NF
14.1%A / <0.01%NF
11.8%A / <0.01%NF
7.2%A / <0.01%NF
7.1%A / <0.01%NF
7.5%A / <0.01%NF
3.8%A / 0.02%NF
3.2%A / 0.03%NF
4.3%A / 0.08%NF
3.2%A / 0.13%NF
3.2%A / 0.26%NF
4.4%A / 0.8%NF
3.3%A / 1.34%NF
6.7%A / 8.92%NF
2.2%A / 8.64%NF
3.1%A / 37.92%NF
0.8%A / 41.84%NF
0
10
20
30
40

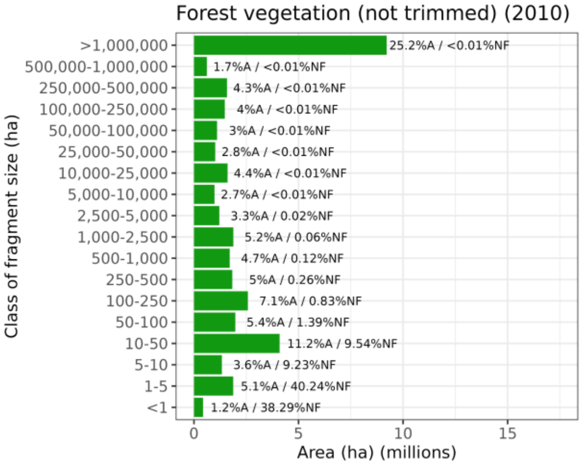
Forest vegetation (not trimmed) (2010)
Class of fragment size (ha)
Area (ha) (millions)
>1,000,000
500,000-1,000,000
250,000-500,000
100,000-250,000
50,000-100,000
25,000-50,000
10,000-25,000
5,000-10,000
2,500-5,000
1,000-2,500
500-1,000
250-500
100-250
50-100
10-50
5-10
1-5
<1
25.2%A / <0.01%NF
1.7%A / <0.01%NF
4.3%A / <0.01%NF
4%A / <0.01%NF
3%A / <0.01%NF
2.8%A / <0.01%NF
4.4%A / <0.01%NF
2.7%A / <0.01%NF
3.3%A / 0.02%NF
5.2%A / 0.06%NF
4.7%A / 0.12%NF
5%A / 0.26%NF
7.1%A / 0.83%NF
5.4%A / 1.39%NF
11.2%A / 9.54%NF
3.6%A / 9.23%NF
5.1%A / 40.24%NF
1.2%A / 38.29%NF
0
5
10
15

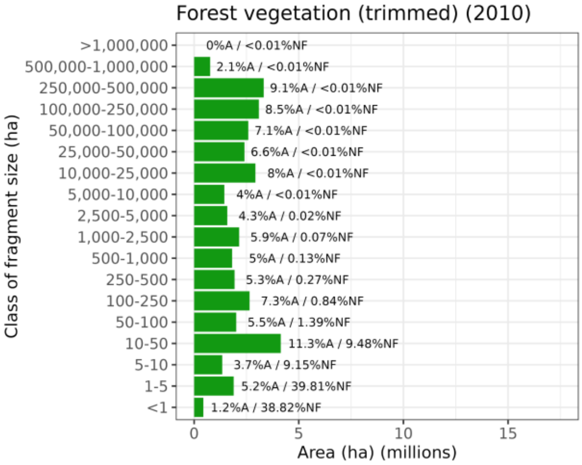
Forest vegetation (trimmed) (2010)
Class of fragment size (ha)
Area (ha) (millions)
>1,000,000
500,000-1,000,000
250,000-500,000
100,000-250,000
50,000-100,000
25,000-50,000
10,000-25,000
5,000-10,000
2,500-5,000
1,000-2,500
500-1,000
250-500
100-250
50-100
10-50
5-10
1-5
<1
0%A / <0.01%NF
2.1%A / <0.01%NF
9.1%A / <0.01%NF
8.5%A / <0.01%NF
7.1%A / <0.01%NF
6.6%A / <0.01%NF
8%A / <0.01%NF
4%A / <0.01%NF
4.3%A / 0.02%NF
5.9%A / 0.07%NF
5%A / 0.13%NF
5.3%A / 0.27%NF
7.3%A / 0.84%NF
5.5%A / 1.39%NF
11.3%A / 9.48%NF
3.7%A / 9.15%NF
5.2%A / 39.81%NF
1.2%A / 38.82%NF
0
5
10
15

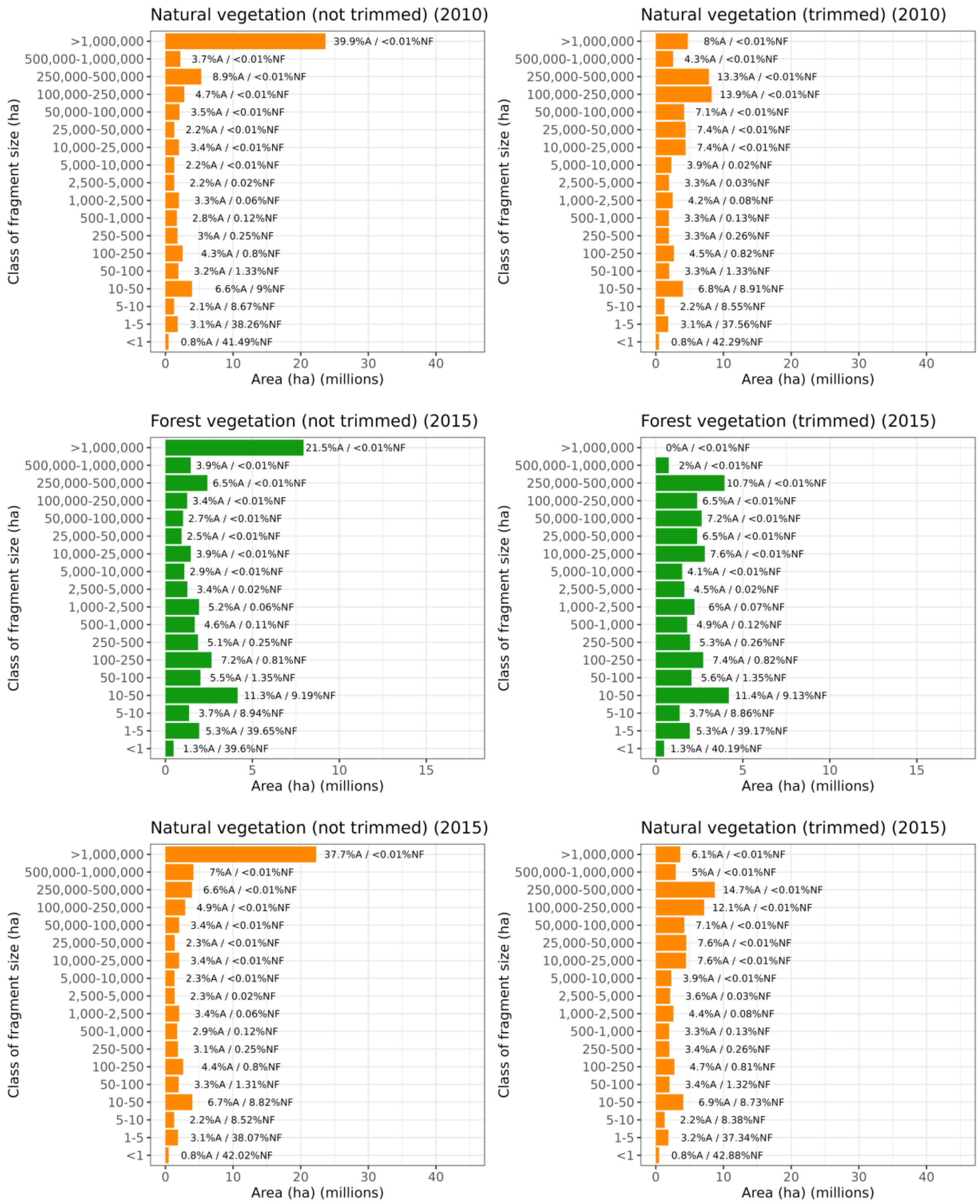


**Figure S3.** Size distribution of forest vegetation and natural vegetation fragments in the Atlantic Forest (1990-2015), without and with road and railways effect. %A: percentage of total area; %NP: percentage of number of fragments.

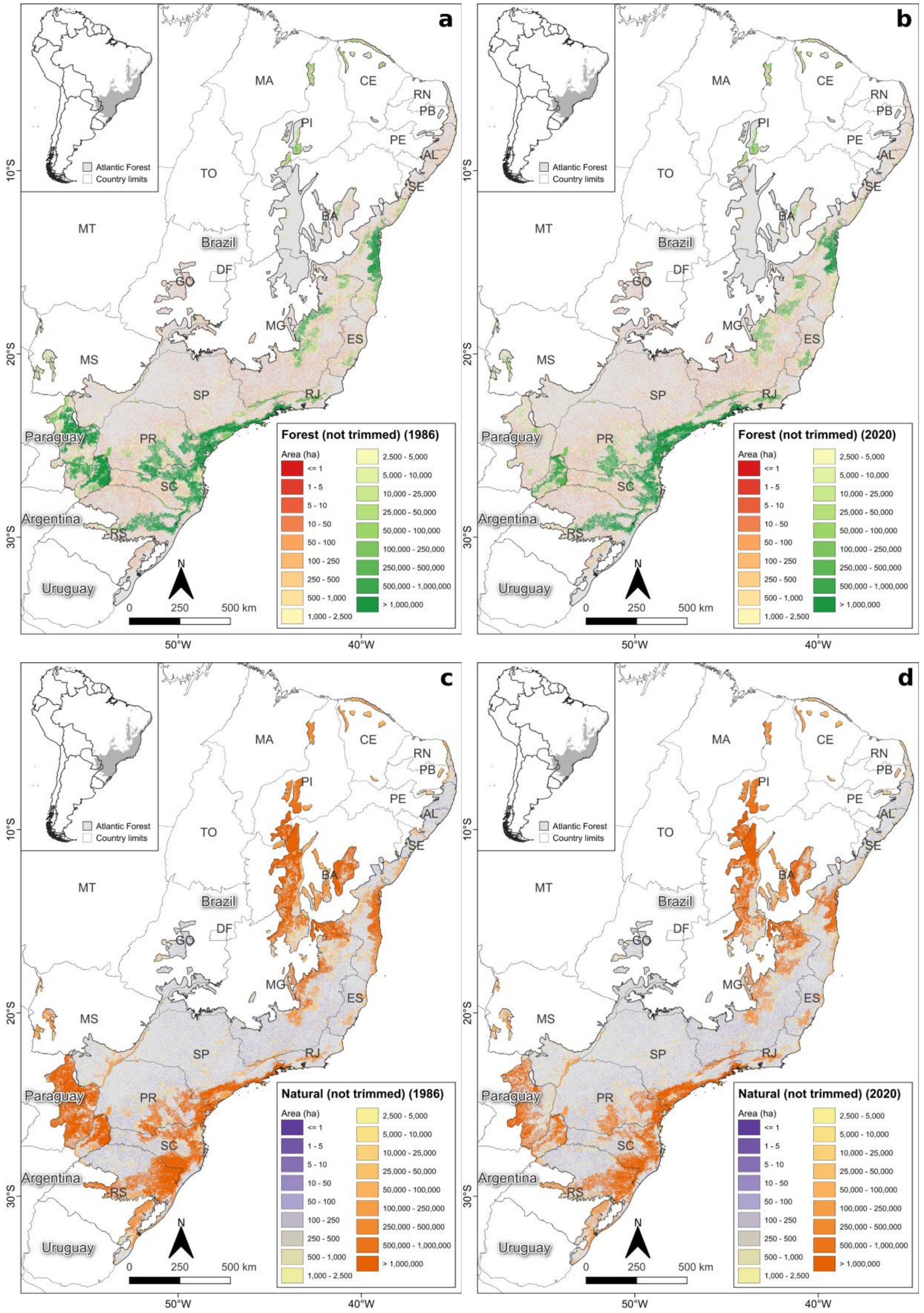


**Figure S4**. Fragment area for forest vegetation in 1986 (a) and 2020 (b), and for natural vegetation in 1986 (c) and 2020 (d), not trimmed for the entire AF.

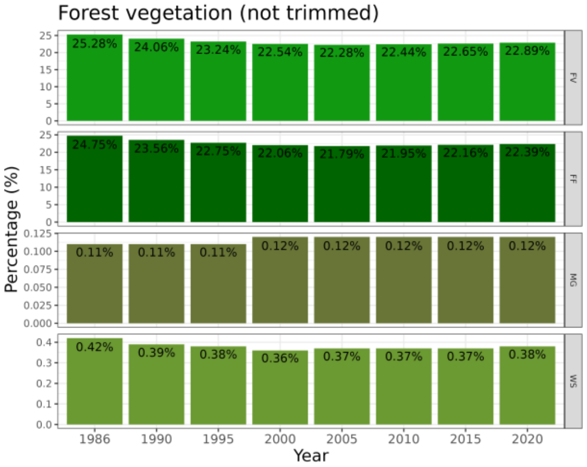
Forest vegetation (not trimmed)
FV: 25.28% 24.06% 23.24% 22.54% 22.28% 22.44% 22.65% 22.89%
FF: 24.75% 23.56% 22.75% 22.06% 21.79% 21.95% 22.16% 22.39%
MG: 0.11% 0.11% 0.11% 0.12% 0.12% 0.12% 0.12% 0.12%
WS: 0.42% 0.39% 0.38% 0.36% 0.37% 0.37% 0.37% 0.38%
Percentage (%)
1986 1990 1995 2000 2005 2010 2015 2020
Year

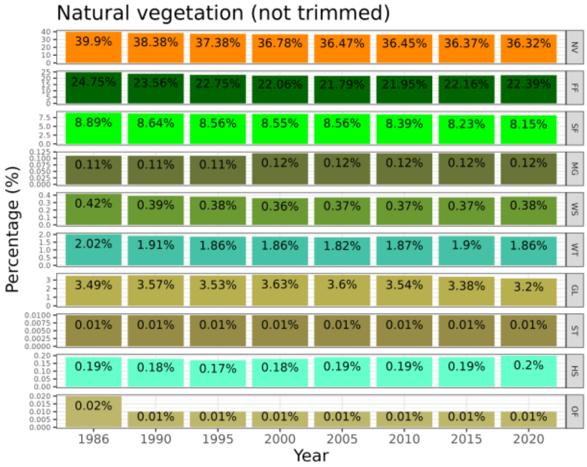
Natural vegetation (not trimmed)
NV: 39.9% 38.38% 37.38% 36.78% 36.47% 36.45% 36.37% 36.32%
FF: 24.75% 23.56% 22.75% 22.06% 21.79% 21.95% 22.16% 22.39%
SF: 8.89% 8.64% 8.56% 8.55% 8.56% 8.39% 8.23% 8.15%
MG: 0.11% 0.11% 0.11% 0.12% 0.12% 0.12% 0.12% 0.12%
WS: 0.42% 0.39% 0.38% 0.36% 0.37% 0.37% 0.37% 0.38%
WT: 2.02% 1.91% 1.86% 1.86% 1.82% 1.87% 1.9% 1.86%
GL: 3.49% 3.57% 3.53% 3.63% 3.6% 3.54% 3.38% 3.2%
ST: 0.01% 0.01% 0.01% 0.01% 0.01% 0.01% 0.01% 0.01%
HS: 0.19% 0.18% 0.17% 0.18% 0.19% 0.19% 0.19% 0.2%
OF: 0.02% 0.01% 0.01% 0.01% 0.01% 0.01% 0.01% 0.01%
Percentage (%)
1986 1990 1995 2000 2005 2010 2015 2020
Year

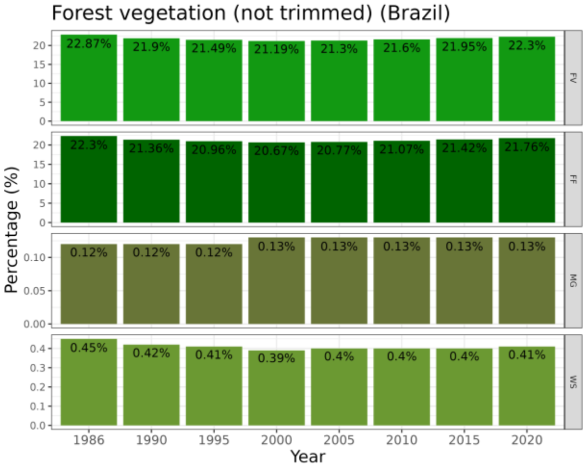
Forest vegetation (not trimmed) (Brazil)
FV: 22.87% 21.9% 21.49% 21.19% 21.3% 21.6% 21.95% 22.3%
FF: 22.3% 21.36% 20.96% 20.67% 20.77% 21.07% 21.42% 21.76%
MG: 0.12% 0.12% 0.12% 0.13% 0.13% 0.13% 0.13% 0.13%
WS: 0.45% 0.42% 0.41% 0.39% 0.4% 0.4% 0.4% 0.41%
Percentage (%)
1986 1990 1995 2000 2005 2010 2015 2020
Year

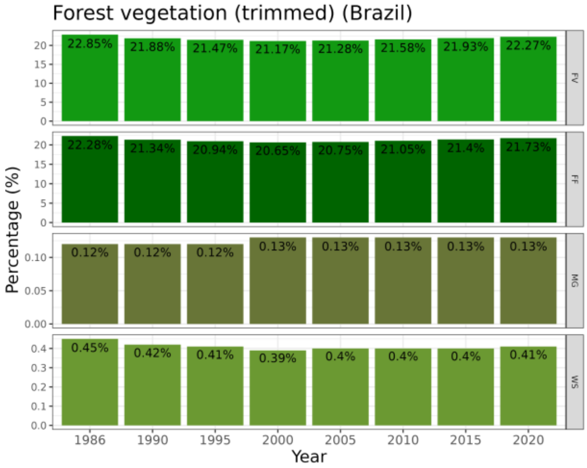
Forest vegetation (trimmed) (Brazil)
FV: 22.85% 21.88% 21.47% 21.17% 21.28% 21.58% 21.93% 22.27%
FF: 22.28% 21.34% 20.94% 20.65% 20.75% 21.05% 21.4% 21.73%
MG: 0.12% 0.12% 0.12% 0.13% 0.13% 0.13% 0.13% 0.13%
WS: 0.45% 0.42% 0.41% 0.39% 0.4% 0.4% 0.4% 0.41%
Percentage (%)
1986 1990 1995 2000 2005 2010 2015 2020
Year

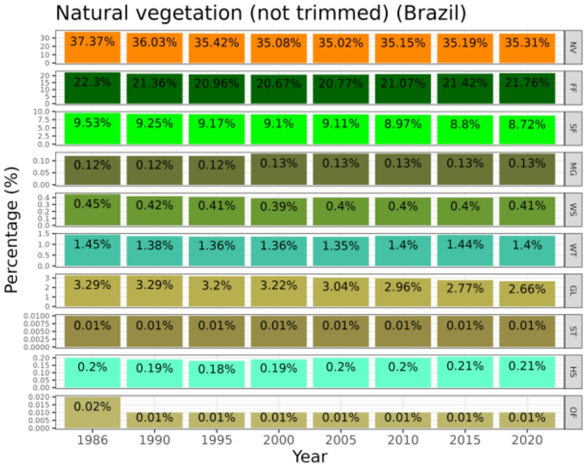
Natural vegetation (not trimmed) (Brazil)
NV: 37.37% 36.03% 35.42% 35.08% 35.02% 35.15% 35.19% 35.31%
FF: 22.3% 21.36% 20.96% 20.67% 20.77% 21.07% 21.42% 21.76%
SF: 9.53% 9.25% 9.17% 9.1% 9.11% 8.97% 8.8% 8.72%
MG: 0.12% 0.12% 0.12% 0.13% 0.13% 0.13% 0.13% 0.13%
WS: 0.45% 0.42% 0.41% 0.39% 0.4% 0.4% 0.4% 0.41%
WT: 1.45% 1.38% 1.36% 1.36% 1.35% 1.4% 1.44% 1.4%
GL: 3.29% 3.29% 3.2% 3.22% 3.04% 2.96% 2.77% 2.66%
ST: 0.01% 0.01% 0.01% 0.01% 0.01% 0.01% 0.01% 0.01%
HS: 0.2% 0.19% 0.18% 0.19% 0.2% 0.2% 0.21% 0.21%
OF: 0.02% 0.01% 0.01% 0.01% 0.01% 0.01% 0.01% 0.01%
Percentage (%)
1986 1990 1995 2000 2005 2010 2015 2020
Year

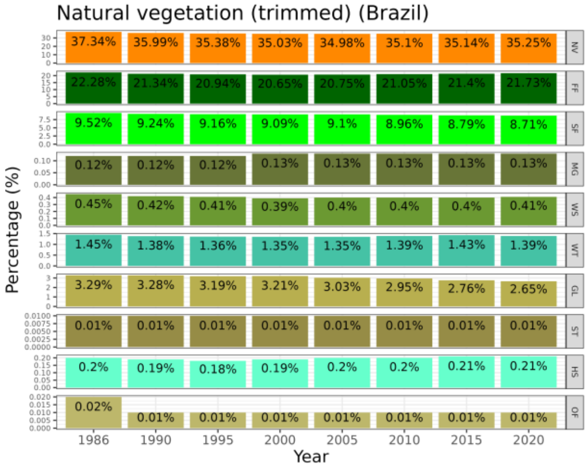
Natural vegetation (trimmed) (Brazil)
NV: 37.34% 35.99% 35.38% 35.03% 34.98% 35.1% 35.14% 35.25%
FF: 22.28% 21.34% 20.94% 20.65% 20.75% 21.05% 21.4% 21.73%
SF: 9.52% 9.24% 9.16% 9.09% 9.1% 8.96% 8.79% 8.71%
MG: 0.12% 0.12% 0.12% 0.13% 0.13% 0.13% 0.13% 0.13%
WS: 0.45% 0.42% 0.41% 0.39% 0.4% 0.4% 0.4% 0.41%
WT: 1.45% 1.38% 1.36% 1.35% 1.35% 1.39% 1.43% 1.39%
GL: 3.29% 3.28% 3.19% 3.21% 3.03% 2.95% 2.76% 2.65%
ST: 0.01% 0.01% 0.01% 0.01% 0.01% 0.01% 0.01% 0.01%
HS: 0.2% 0.19% 0.18% 0.19% 0.2% 0.2% 0.21% 0.21%
OF: 0.02% 0.01% 0.01% 0.01% 0.01% 0.01% 0.01% 0.01%
Percentage (%)
1986 1990 1995 2000 2005 2010 2015 2020
Year

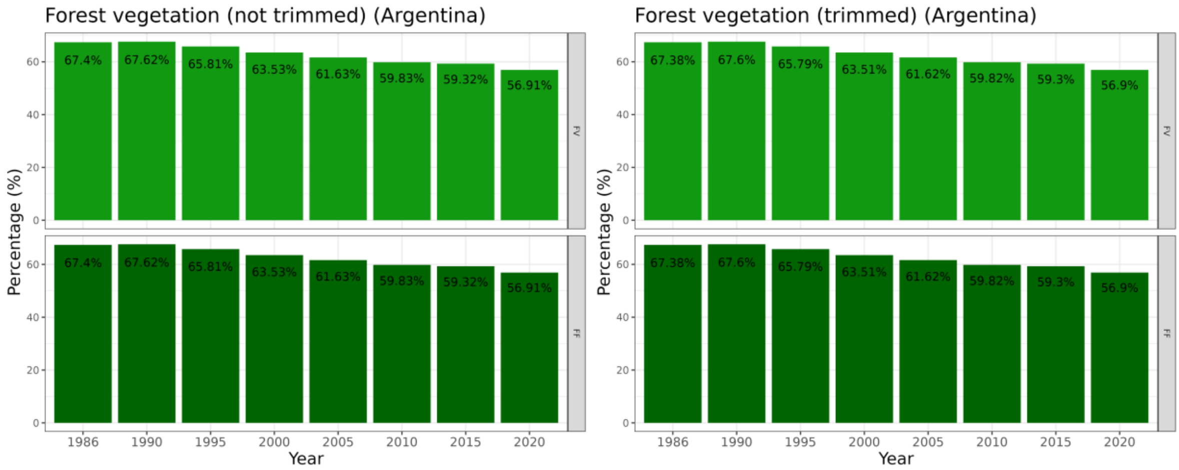

Forest vegetation (not trimmed) (Argentina)
Forest vegetation (trimmed) (Argentina)
Percentage (%)
Year
1986 1990 1995 2000 2005 2010 2015 2020
FV
FF
67.4% 67.62% 65.81% 63.53% 61.63% 59.83% 59.32% 56.91%
67.38% 67.6% 65.79% 63.51% 61.62% 59.82% 59.3% 56.9%


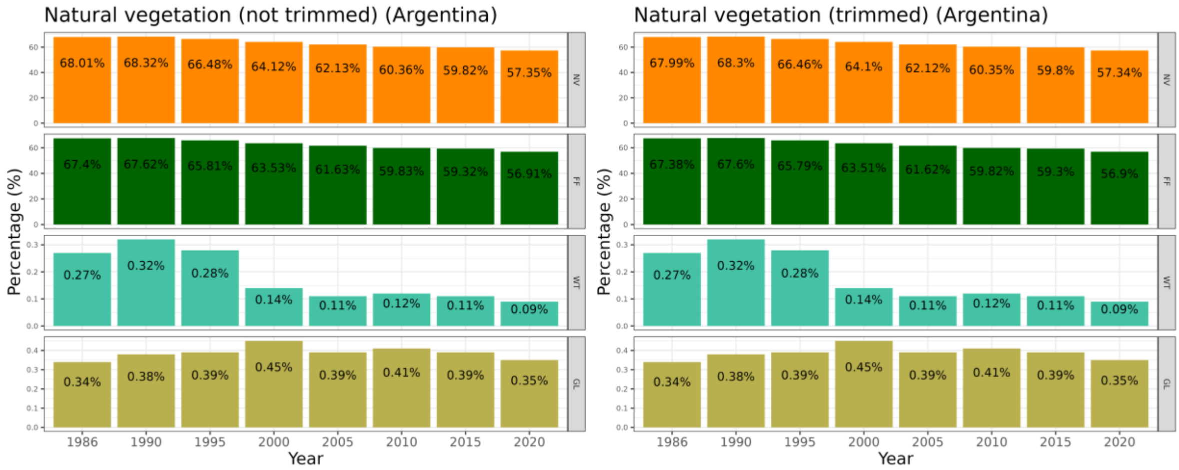

Natural vegetation (not trimmed) (Argentina)
Natural vegetation (trimmed) (Argentina)
Percentage (%)
Year
1986 1990 1995 2000 2005 2010 2015 2020
NV
FF
WT
GL
68.01% 68.32% 66.48% 64.12% 62.13% 60.36% 59.82% 57.35%
67.4% 67.62% 65.81% 63.53% 61.63% 59.83% 59.32% 56.91%
0.27% 0.32% 0.28% 0.14% 0.11% 0.12% 0.11% 0.09%
0.34% 0.38% 0.39% 0.45% 0.39% 0.41% 0.39% 0.35%
67.99% 68.3% 66.46% 64.1% 62.12% 60.35% 59.8% 57.34%
67.38% 67.6% 65.79% 63.51% 61.62% 59.82% 59.3% 56.9%

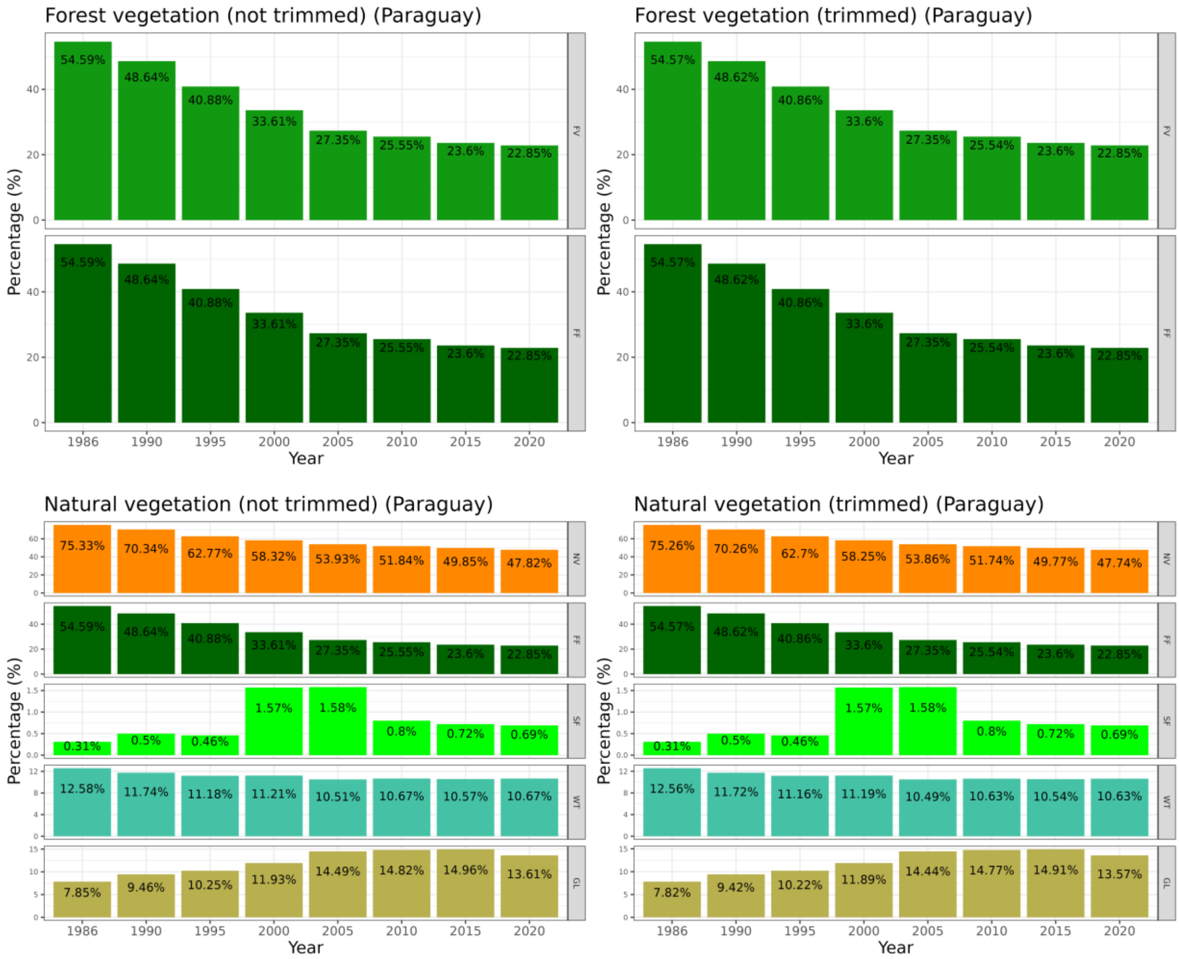


**Figure S5**. FV and NV cover per year for the entire Atlantic Forest, Brazil, Argentina, and Paraguay, trimmed and not trimmed. Abbreviations for FV cover are: FV = Forest vegetation, FF = Forest formation, MG = Mangrove, WS = Wooded sandbank vegetation (*restinga*). Abbreviations for NV cover are: NV = Natural vegetation, FF = Forest formation, SF = Savanna formation, MG = Mangrove, WS = Wooded sandbank vegetation (*restinga*), WT = Wetland, GL = Grassland, ST = Salt flat, HS = Herbaceous sandbank vegetation, OF = Other non-forest formations. Class colors follow MapBiomas collection 7 and MapBiomas Bosque Atlántico collection 2 legends.

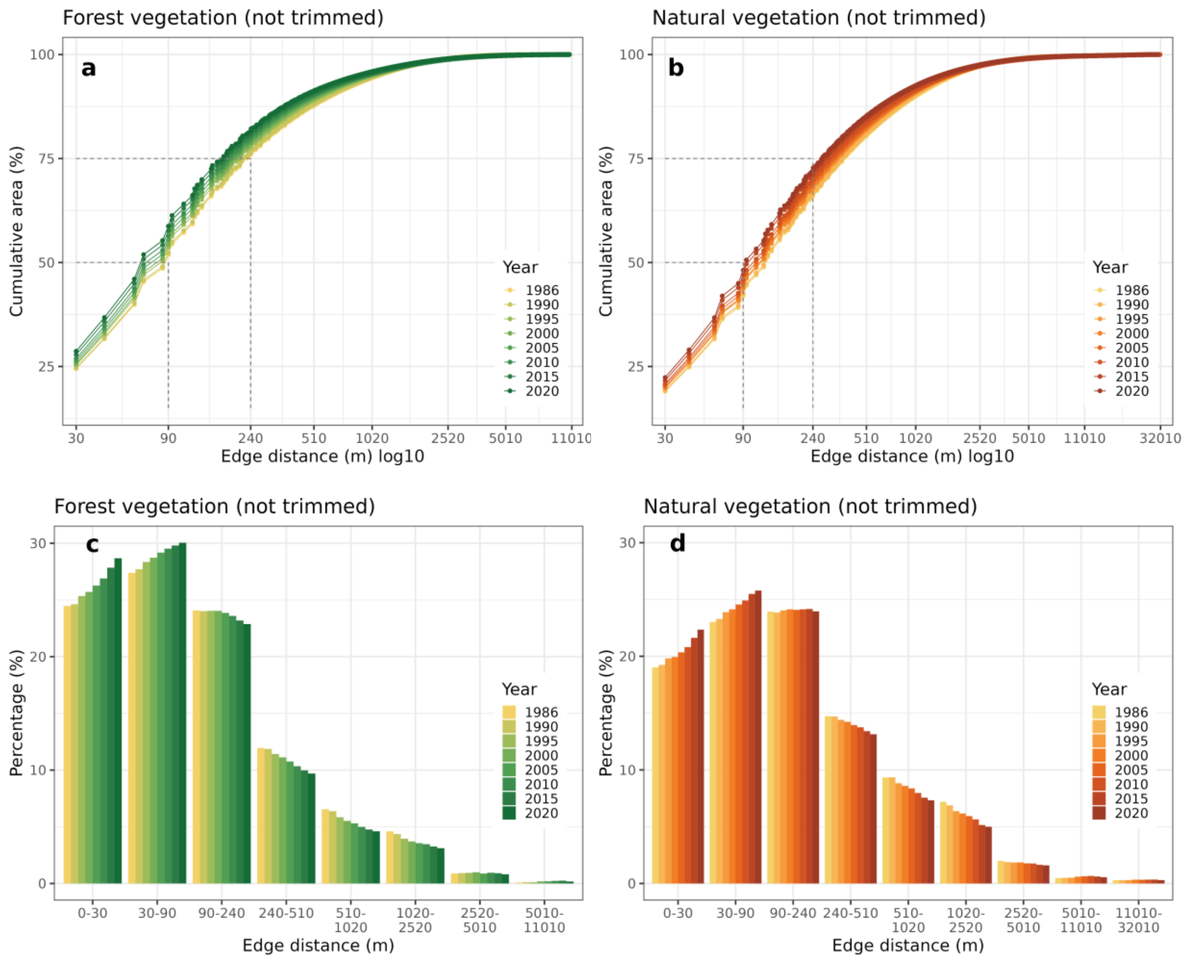


**Figure S6.** Cumulative (a-b) and per class (c-d) area under edge effect at different edge depths for the FV and NV remaining in Atlantic Forest not trimmed.

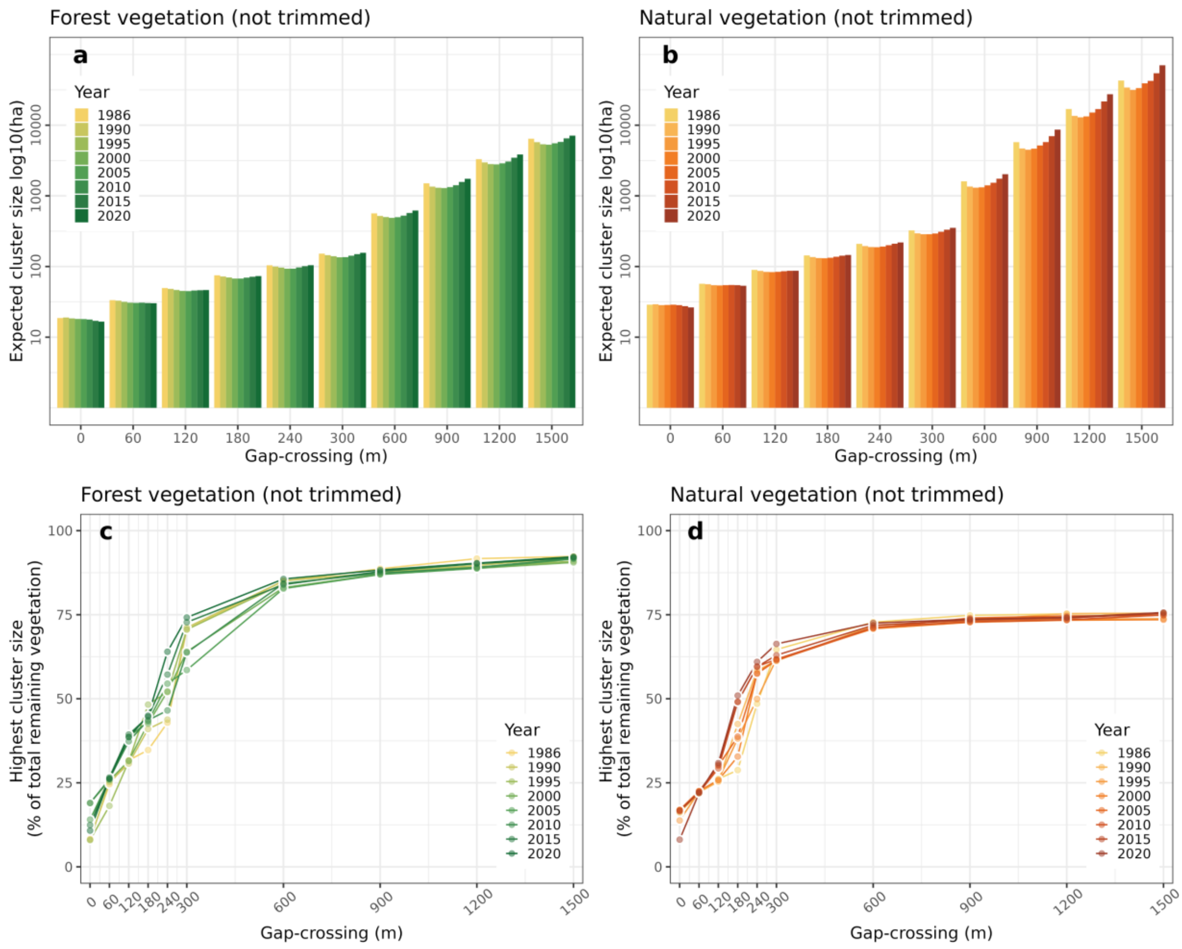


**Figure S7.** Expected cluster size (a-b) (average functional size; ha) of functionally connected fragments of FV and NV for different functional distances values (meters) for the Atlantic Forest. Highest functionally connected forest cluster (c-d) (% of total remaining of FV and NV) estimated across varying functional distances (m), for the Atlantic Forest not trimmed by roads and railways.

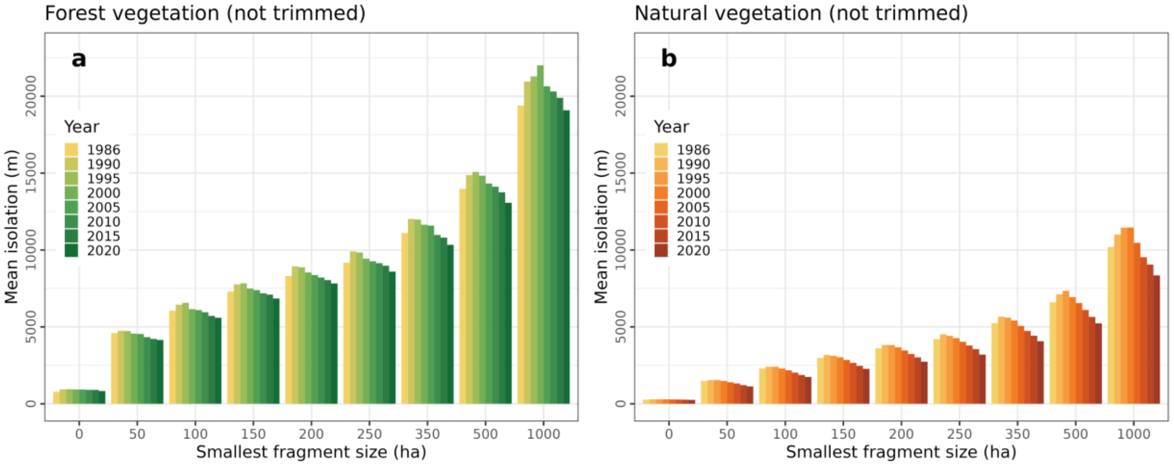


**Figure S8**. Influence of the smallest fragment size (ha) on the mean isolation (m) between fragments of FV and NV for the Atlantic Forest: mean isolation (a-b) not trimmed. Smallest fragments size: 0 ha (all fragments), 50 ha, 100 ha, 150 ha, 200 ha, 250 ha, 350 ha, 500 ha, and 1000 ha.

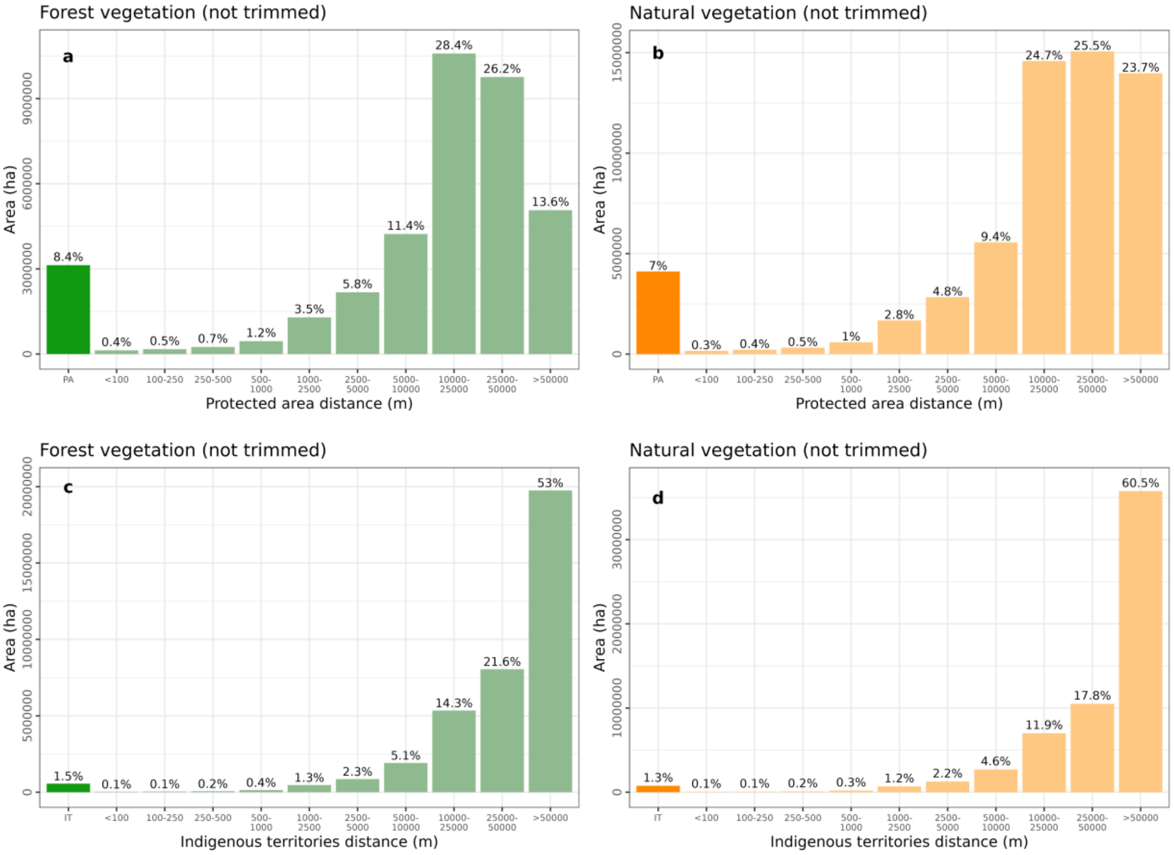


**Figure S9.** Remaining vegetation within (area and percentage) and its distance by class (meters) from protected areas (PA; a – FV and b – NV) and indigenous lands (IT; c – FV and d – NV) to AF not trimmed.

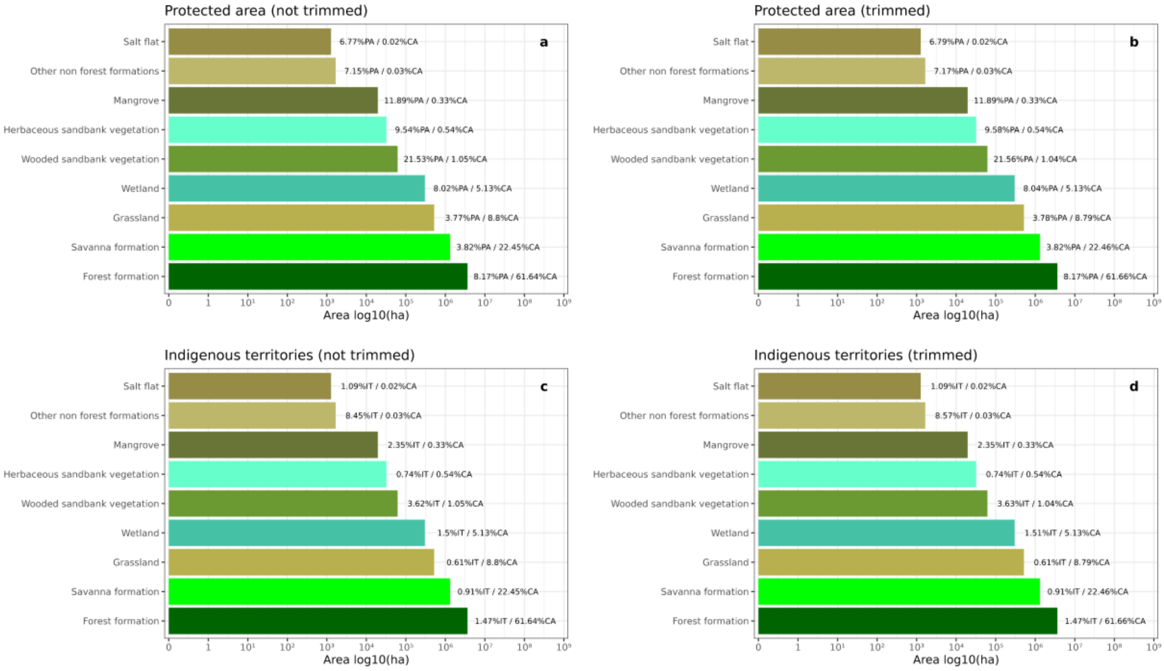


**Figure S10**. Remaining classes cover for FV and NV (area and percentage) within protected areas (PA) (a-b) and indigenous territories (TI) (c-d) for the Atlantic Forest not trimmed and trimmed. %PA: percentage of class area inside PA in relation to the total area of the class; %IT: percentage of class area inside IT in relation to the total area of the class; %CA: percentage of the class area in relation to the total area of vegetation for PA and IT.

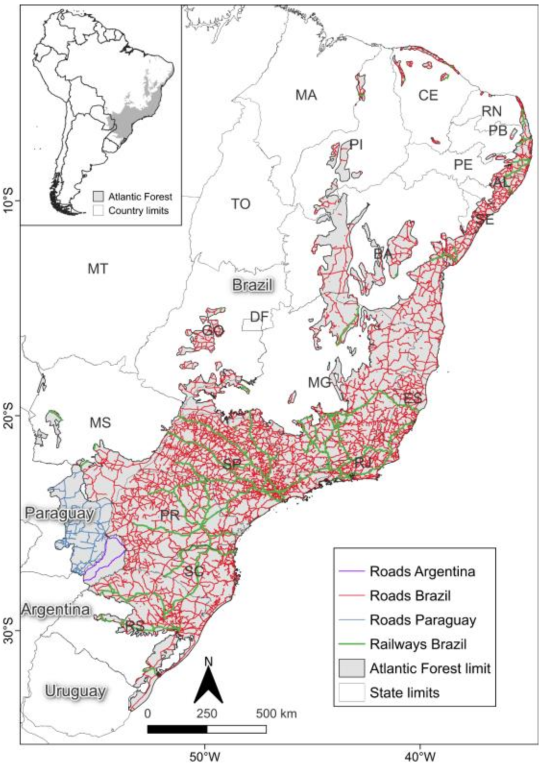


**Figure S11.** Roads and railways used to trim the FV and NV fragments.

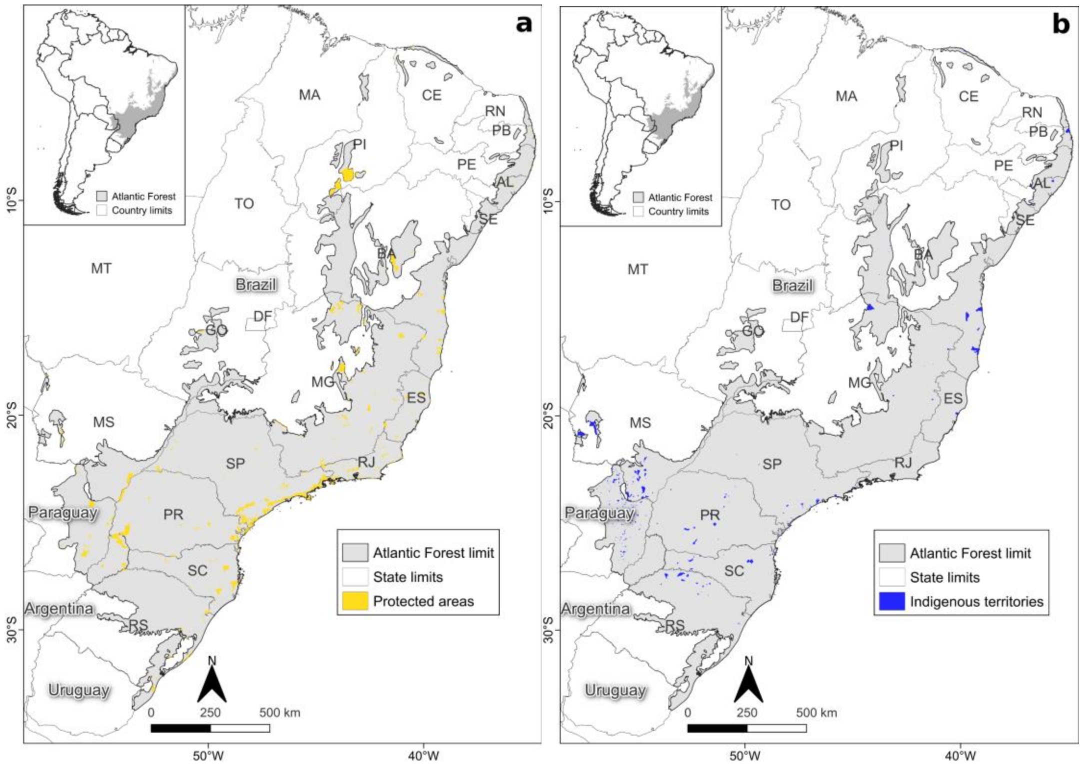


**Figure S12.** Protected areas (a) and indigenous territories (b) within the Atlantic Forests of South America.

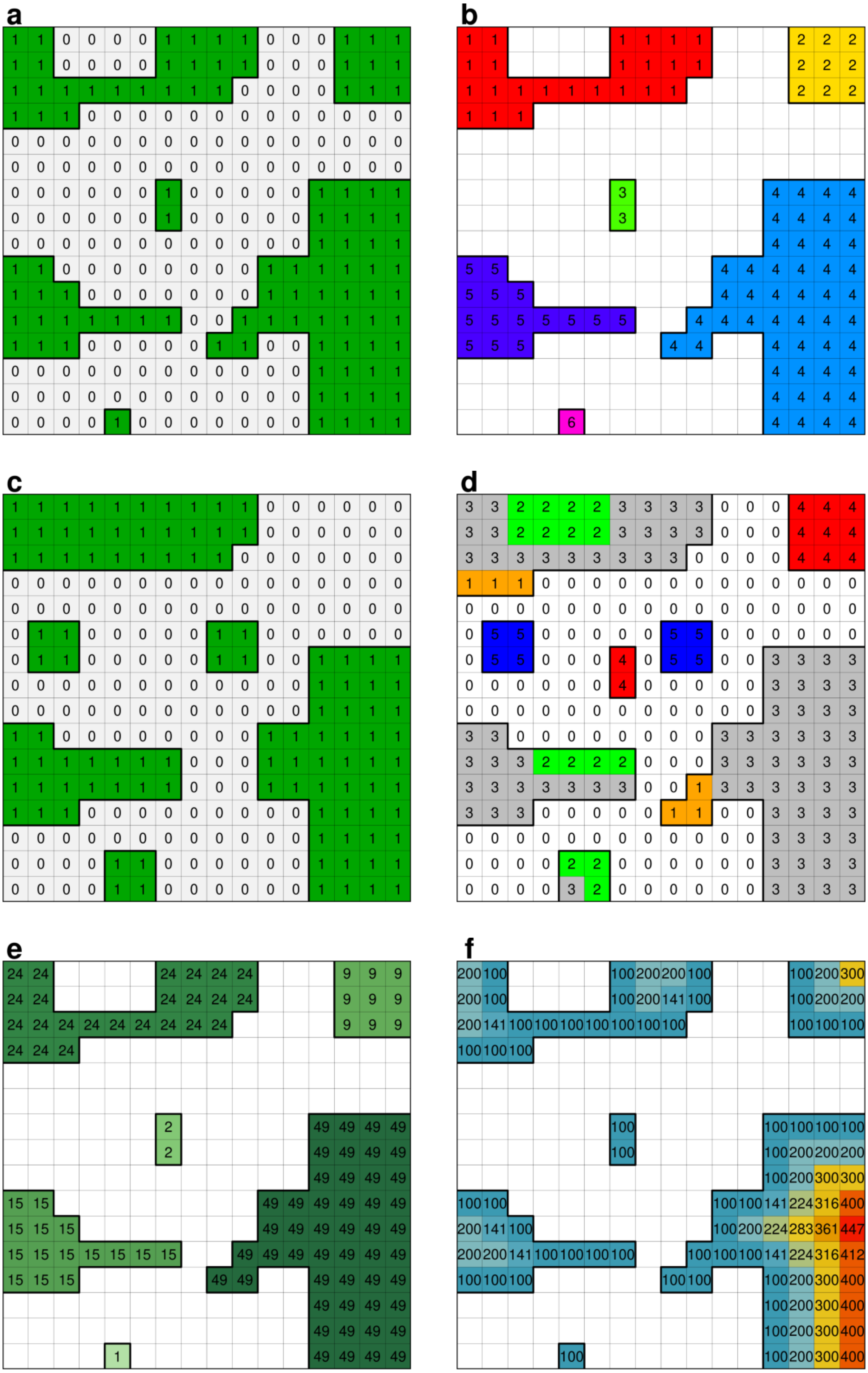
a
b
c
d
e
f

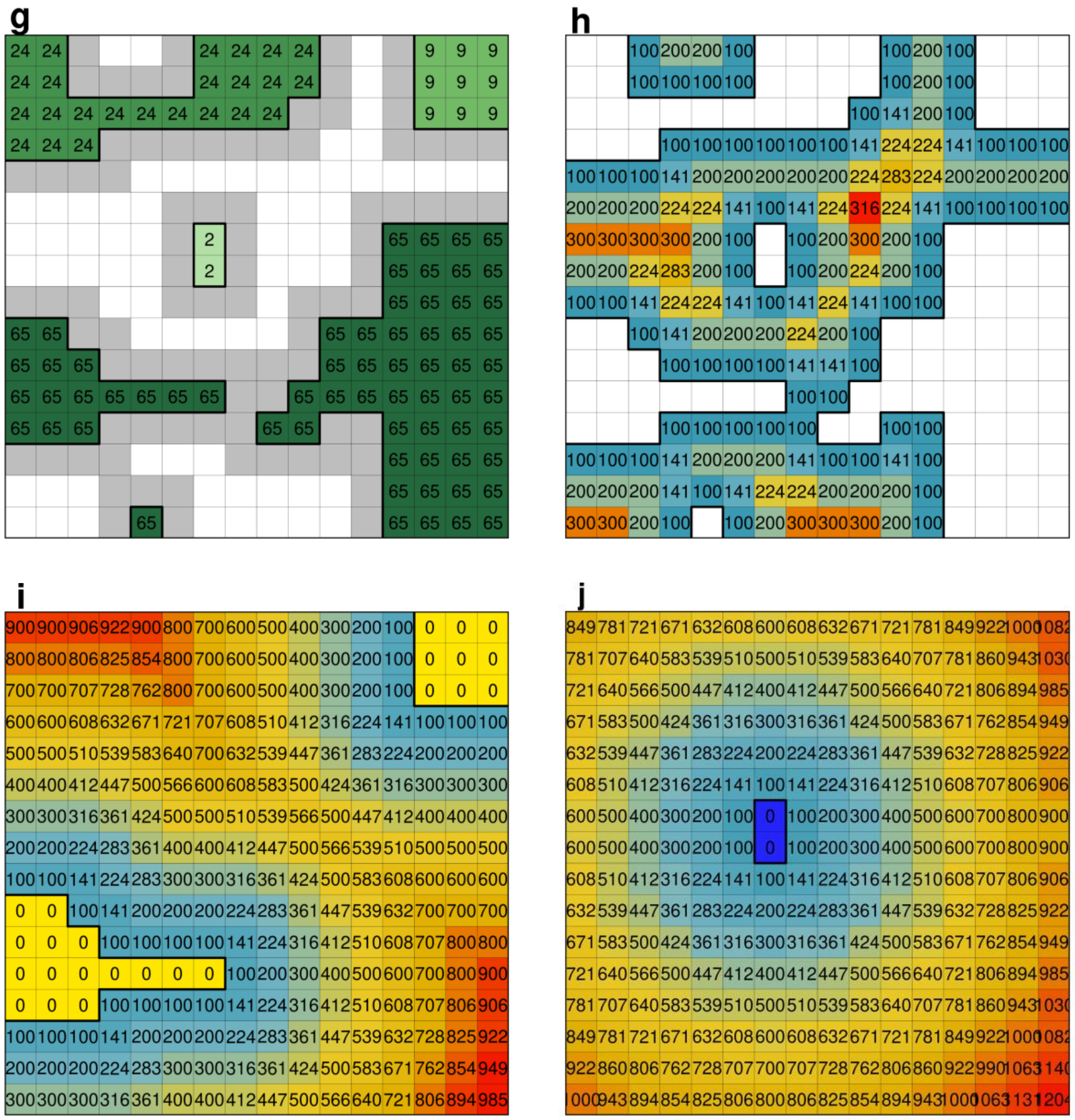


**Figure S13**. Illustration exemplifying the landscape metrics used to analyze the landscape structure of the AF. Here, the rasters have 100 m of spatial resolution. (a) toy landscape containing binary forest and non-forest data, (b) number of fragments; (c) second toy landscape; (d) temporal fragment analysis—matrix (0), loss area (1), gain area (2), stable area (3), loss fragment (4), gain fragment (5); (e) fragment size; (f) edge area; (g) functional connectivity; (h) isolation; (i) distance from protected areas; and (j) distance from indigenous territories.

### Tables

**Table S1**. Atlantic Forest (AF) delimitations used to compose the AF "integrated delimitation" adapted from Muylaert et al. (2018).

| Reference | Area (ha) | Percentage in relation to Muylaert et al. (2018) | Figure |
|---|---|---|---|
| Muylaert et al. (2018) | 162,742,129 | 100.0% | Fig. S1a |
| Muylaert et al. (2018) – Brazil | 151,470,253 | 93.1% | Fig. S1a |
| Muylaert et al. (2018) – Argentina | 2,668,855 | 1.64% | Fig. S1a |
| Muylaert et al. (2018) – Paraguay | 8,603,022 | 5.29% | Fig. S1a |
| Atlantic Forest Law (2006) | 128,779,574 | 79.1% | Fig. S1b |
| Da Silva and Casteleti (2003) | 136,422,859 | 83.8% | Fig. S1c |
| IBGE (2004) | 111,751,193 | 68.7% | Fig. S1d |
| IBGE (2019) | 110,656,961 | 68.0% | Fig. S1e |
| Dinerstein et al. (2017) | 120,492,216 | 74.0% | Fig. S1f |

**Table S2**. Land use and land cover and MapBiomas class code used to compose the "Forest Vegetation" (FV) and "Natural Vegetation" (NV).

| Vegetation class | Land use and land cover class | Land use and land cover class abbreviation | MapBiomas class code |
|---|---|---|---|
| Forest Vegetation (FV) | Forest Formation | FF | 3 |
| | Mangrove | MG | 5 |
| | Wooded Sandbank Vegetation (*restinga*) | WS | 49 |
| Natural Vegetation (NV) | Forest Formation | FF | 3 |
| | Savanna Formation | SF | 4 |
| | Mangrove | MG | 5 |
| | Wooded Sandbank Vegetation (*restinga*) | WS | 49 |
| | Wetland | WT | 11 |
| | Grassland | GL | 12 |
| | Salt Flat | ST | 32 |
| | Herbaceous Sandbank Vegetation | HS | 50 |
| | Other Non-forest Formations | OF | 13 |

**Table S3**. Landscape and topographic metrics used to analyze the structure of the landscapes of the AF.

| Metric | Description | Class |
|---|---|---|
| Number of fragments and fragment size. | Number of fragments, fragment size and percentage of habitat cover for different size classes. | fragment size classes (ha): <1, 1–5, 5–10, 10–50, 50–100, 100–250, 250–500, 500–1000, 1000–2500, 2500–5000, 5000–10000, 10000–25000, 25000–50000, 50000–100000, 100000–250000, 250000–500000, 500000–1000000, and >1000000. |
| Temporal dynamics of the landscape: area and number of fragments. | Areas of increase, reduction, and stability of fragments that remained throughout time, and area and number of fragments that disappeared and appeared. | Values (Fig. S13): accounting area of loss (1), gain (2), and stable (3) of the stains that remained; number and area of fragments that disappeared (4) and appeared (5). |
| Edge area. | Percentage of habitat area submitted to edge effects for different edge widths. | Edge widths (m) (pixel size): <30, 30–90, 90–240, 240–510, 510–1020, 1020–2520, 2520–5010, 5010–11010, and 11010–32010. |
| Functional connectivity. | Area of functionally connected fragments, considering different distance rules for fragment linkage. | Gap-crossing (m) (pixel size): 0, 60, 120, 180, 240, 300, 600, 900, 1200, and 1500. |
| Mean isolation. | Mean isolation to the nearest habitat fragment. To analyze the effect of small fragments in estimating isolation, the smallest fragments were successively removed. | Size of the small fragments removed (ha): 0 (i.e., no fragments removed), <50, <100, <150, <200, <250, <350, <500, and <1000. |
| Distance from Protected Areas and Indigenous Territories. | Distance of any given habitat pixel to the nearest Protected Area and Indigenous Territories. | Distance classes (m): 0 (i.e., inside a Protected Area or Indigenous Territories), <100, 100–250, 250–500, 500–1000, 1000–2500, 2500–5000, 5000–10000, 10000–25000, 25000–50000, and >50000. |

**Table S4**. Remaining FV and NV for the entire Atlantic Forest between 1986 and 2020, trimmed and not trimmed. For each year and vegetation scenario, the following values are presented: total percentage, number of fragments, and descriptive statistics (average, standard deviation, median and maximum) in hectares.

| Year | Scenario | Total percentage (%) | Number of fragments | Total area (ha) | Average area (ha) | Standard deviation area (ha) | Median area (ha) | Maximum area (ha) |
|---|---|---|---|---|---|---|---|---|
| 1986 | FV not trimmed | 25.28 | 2,201,796 | 41,132,358 | 18.68 | 3,687 | 1.35 | 3,397,448 |
| 1986 | FV trimmed | 25.26 | 2,230,509 | 41,096,833 | 18.42 | 1,296 | 1.35 | 830,496 |
| 1986 | NV not trimmed | 39.9 | 2,251,224 | 64,909,217 | 28.83 | 9,245 | 1.17 | 10,490,715 |
| 1986 | NV trimmed | 39.86 | 2,301,140 | 64,832,991 | 28.17 | 2,956 | 1.17 | 2,711,898 |
| 1990 | FV not trimmed | 24.06 | 2,059,724 | 39,164,420 | 19.01 | 3,472 | 1.44 | 3,158,986 |
| 1990 | FV trimmed | 24.04 | 2,086,758 | 39,130,573 | 18.75 | 1,269 | 1.44 | 877,595 |
| 1990 | NV not trimmed | 38.38 | 2,136,171 | 62,471,881 | 29.24 | 9,277 | 1.26 | 10,532,927 |
| 1990 | NV trimmed | 38.34 | 2,185,346 | 62,397,614 | 28.55 | 2,953 | 1.26 | 2,677,168 |
| 1995 | FV not trimmed | 23.24 | 2,051,750 | 37,825,688 | 18.44 | 3,336 | 1.44 | 3,007,756 |
| 1995 | FV trimmed | 23.22 | 2,080,013 | 37,791,570 | 18.17 | 1,170 | 1.44 | 803,755 |
| 1995 | NV not trimmed | 37.38 | 2,137,418 | 60,831,540 | 28.46 | 7,985 | 1.26 | 8,386,742 |
| 1995 | NV trimmed | 37.34 | 2,188,098 | 60,757,173 | 27.77 | 2,893 | 1.26 | 2,714,043 |
| 2000 | FV not trimmed | 22.54 | 2,024,772 | 36,688,790 | 18.12 | 5,092 | 1.44 | 6,959,510 |
| 2000 | FV trimmed | 22.52 | 2,052,621 | 36,655,360 | 17.86 | 1,150 | 1.44 | 770,101 |
| 2000 | NV not trimmed | 36.78 | 2,091,472 | 59,861,952 | 28.62 | 8,774 | 1.35 | 10,157,618 |
| 2000 | NV trimmed | 36.74 | 2,143,016 | 59,786,308 | 27.9 | 2,810 | 1.35 | 2,662,722 |
| 2005 | FV not trimmed | 22.28 | 2,009,506 | 36,260,960 | 18.04 | 4,020 | 1.53 | 5,095,019 |
| 2005 | FV trimmed | 22.26 | 2,039,272 | 36,225,866 | 17.76 | 1,132 | 1.44 | 806,599 |
| 2005 | NV not trimmed | 36.47 | 2,060,924 | 59,344,869 | 28.8 | 8,663 | 1.35 | 10,017,365 |

| | | | | | | | | |
|---|---|---|---|---|---|---|---|---|
| 2005 | NV trimmed | 36.42 | 2,114,253 | 59,266,718 | 28.03 | 2,764 | 1.35 | 2,603,116 |
| 2010 | FV not trimmed | 22.44 | 2,052,341 | 36,522,265 | 17.8 | 5,034 | 1.44 | 6,937,266 |
| 2010 | FV trimmed | 22.41 | 2,084,891 | 36,484,973 | 17.5 | 1,100 | 1.44 | 763,970 |
| 2010 | NV not trimmed | 36.45 | 2,085,299 | 59,314,001 | 28.44 | 8,356 | 1.35 | 9,792,496 |
| 2010 | NV trimmed | 36.39 | 2,142,699 | 59,232,818 | 27.64 | 2,700 | 1.35 | 2,625,471 |
| 2015 | FV not trimmed | 22.65 | 2,161,797 | 36,867,412 | 17.05 | 3,705 | 1.44 | 4,582,605 |
| 2015 | FV trimmed | 22.62 | 2,199,425 | 36,826,511 | 16.74 | 1,081 | 1.35 | 744,569 |
| 2015 | NV not trimmed | 36.37 | 2,151,325 | 59,180,724 | 27.51 | 8,195 | 1.35 | 9,895,304 |
| 2015 | NV trimmed | 36.3 | 2,215,181 | 59,096,027 | 26.68 | 2,574 | 1.26 | 2,532,163 |
| 2020 | FV not trimmed | 22.89 | 2,244,015 | 37,251,477 | 16.6 | 3,343 | 1.35 | 4,010,367 |
| 2020 | FV trimmed | 22.86 | 2,287,893 | 37,206,313 | 16.26 | 1,026 | 1.35 | 698,736 |
| 2020 | NV not trimmed | 36.32 | 2,241,089 | 59,110,442 | 26.38 | 6,103 | 1.26 | 4,779,875 |
| 2020 | NV trimmed | 36.27 | 2,310,919 | 59,022,773 | 25.54 | 2,406 | 1.26 | 2,120,105 |

**Table S5**. Dynamics of the landscape for the entire Atlantic Forest, from 1986-2005 and from 2005-2020 for two scenarios: not trimmed and trimmed. Balance of total area represents the subtraction of area gained minus area lost. Balance of total area (%) is in terms of percentage in relation to the total AF area. Balance of number of fragments represents the subtraction of the number of fragments gained minus the number of fragments lost. Mean size of fragments gained (ha), and mean size of fragments lost (ha) represent the average size of fragments gained and lost. The table with all measures is presented in Table S3.

| Period | Scenario | Balance of total area (ha) | Balance of total area (%) | Balance of number of fragments | Mean size of fragments gained (ha) | Mean size of fragments lost (ha) |
|---|---|---|---|---|---|---|
| 1986-2005 | FV not trimmed | -4,871,398 | -2.99 | -242,715 | 1.14 | 1.35 |
| 1986-2005 | FV trimmed | -4,870,967 | -2.99 | -242,678 | 1.12 | 1.33 |
| 1986-2005 | NV not trimmed | -5,564,348 | -3.42 | -227,621 | 1.11 | 1.21 |
| 1986-2005 | NV trimmed | -5,566,273 | -3.42 | -226,734 | 1.08 | 1.20 |
| 2005-2020 | FV not trimmed | 990,517 | 0.61 | 373,616 | 1.08 | 0.97 |
| 2005-2020 | FV trimmed | 980,447 | 0.60 | 385,798 | 1.06 | 0.96 |
| 2005-2020 | NV not trimmed | -234,427 | -0.14 | 301,911 | 1.06 | 0.96 |
| 2005-2020 | NV trimmed | -243,945 | -0.15 | 314,062 | 1.03 | 0.94 |

**Table S6**. Remaining FV and NV for different AF delimitations for the year of 2020, trimmed and not trimmed by roads and railways. For each delimitation and vegetation scenario are presented: vegetation percentage, number of fragments, and total and average area in hectares. The AF delimitations can be seen in Fig. S1.

| AF delimitation | Scenario | Vegetation percentage (%) | Number of fragments | Total area (ha) | Average area (ha) |
|---|---|---|---|---|---|
| Atlantic Forest Law (2006) | FV not trimmed | 24.07 | 1,788,188 | 30,992,081 | 17.33 |
| Atlantic Forest Law (2006) | FV trimmed | 24.03 | 1,827,380 | 30,951,928 | 16.94 |
| Atlantic Forest Law (2006) | NV not trimmed | 35.98 | 1,835,652 | 46,333,549 | 25.24 |
| Atlantic Forest Law (2006) | NV trimmed | 35.93 | 1,892,400 | 46,264,936 | 24.45 |
| Da Silva and Casteleti (2003) | FV not trimmed | 23.18 | 1,902,365 | 31,622,147 | 16.62 |
| Da Silva and Casteleti (2003) | FV trimmed | 23.15 | 1,944,184 | 31,579,924 | 16.24 |
| Da Silva and Casteleti (2003) | NV not trimmed | 34.36 | 1,933,978 | 46,875,495 | 24.24 |
| Da Silva and Casteleti (2003) | NV trimmed | 34.31 | 1,993,589 | 46,806,306 | 23.48 |
| IBGE (2004) | FV not trimmed | 26.06 | 1,641,173 | 29,122,577 | 17.74 |
| IBGE (2004) | FV trimmed | 26.02 | 1,679,756 | 29,083,232 | 17.31 |
| IBGE (2004) | NV not trimmed | 31.84 | 1,671,550 | 35,586,103 | 21.29 |
| IBGE (2004) | NV trimmed | 31.79 | 1,722,521 | 35,529,739 | 20.63 |
| IBGE (2019) | FV not trimmed | 26.5 | 1,649,485 | 29,326,354 | 17.78 |
| IBGE (2019) | FV trimmed | 26.47 | 1,688,418 | 29,286,695 | 17.35 |
| IBGE (2019) | NV not trimmed | 31.5 | 1,662,346 | 34,859,089 | 20.97 |
| IBGE (2019) | NV trimmed | 31.45 | 1,711,606 | 34,805,009 | 20.33 |
| Dinerstein et al. (2017) | FV not trimmed | 26.7 | 1,796,063 | 32,173,468 | 17.91 |
| Dinerstein et al. (2017) | FV trimmed | 26.67 | 1,834,102 | 32,133,736 | 17.52 |
| Dinerstein et al. (2017) | NV not trimmed | 33.97 | 1,767,078 | 40,934,679 | 23.17 |
| Dinerstein et al. (2017) | NV trimmed | 33.92 | 1,821,027 | 40,870,573 | 22.44 |

UNEP-WCMC and IUCN, 2022. Protected Planet: The World Database on Protected Areas (WDPA) and World Database on Other Effective Area-based Conservation Measures (WD-OECM) [Online], November 2022, Cambridge, UK: UNEP-WCMC and IUCN. Available at: www.protectedplanet.net.
